# Electrical Coordinated Reset Stimulation Induces Network Desynchronization in an in Vivo Model of Status Epilepticus

**DOI:** 10.1101/2024.11.06.621943

**Authors:** Daniel Ehrens, Fadi Aeed, Yara Otor, Vivek Charu, Babak Razavi, Sridevi V. Sarma, Yitzhak Schiller, Peter A. Tass

## Abstract

Epilepsy, a neurological disorder characterized by recurrent seizures, profoundly impacts individuals worldwide. Various electrical stimulation protocols have been investigated to mitigate epileptic seizures, among which Coordinated Reset (CR) stimulation may have potential for inducing long-lasting neural desynchronization. This study explores the acute effects of CR stimulation on synchronization dynamics during Status Epilepticus (SE) in an in vivo animal model. An electrographically sustained seizure-state was induced via 4-aminopyridine (4AP) administration to CA3. Custom-designed electrode probes were implanted to facilitate simultaneous recording and electrical stimulation. Analytical univariate and bivariate features were constructed from the LFP time-series recording. Feature metrics focused on spike synchronization metrics and continuous signal analysis of amplitude, spectral power and phase synchronization across electrode pairs and frequency bands. Significance of modulation was assessed through permutation testing of the observed differences between the CR-stimulated group (N=5) compared to the control (no stimulation) group (N=3) during SE. Results showed overall decrease in amplitude and power univariate features, and a significant modulation of bivariate synchronization and connectivity measures across the spectrum between the CR stimulation and control group. Our findings underscore the potential effectiveness of CR stimulation in attenuating excessive neural synchronization, paving the way for further exploration of CR stimulation as a viable intervention for network desynchronization of epileptiform activity and subsequently treatment of seizures.

## 1. Introduction

Epilepsy is a chronic neurological disorder characterized by recurrent, unprovoked seizures, impacting approximately 70 million individuals globally [1]. Approximately 30% of individuals with epilepsy are afflicted with Drug-Resistant Epilepsy (DRE), characterized by the persistence of seizures despite with different Anti-Seizure Medication (ASMs) [2–4]. Individuals with DRE may benefit from surgical resection with the aim of surgically removing the epileptogenic zone, a specific region or network identified as crucial for seizure initiation and spread [5,6].

However, surgical resection for DRE often necessitates surgical intervention, which is irreversible, entails substantial risks, and post-operative challenges and outcomes are highly variable [7,8]. In recent years, electrical stimulation has emerged as a promising alternative to surgical resection for DRE treatment [9–11], offering therapeutic options such as Deep Brain Stimulation (DBS) [12], Vagus Nerve Stimulation (VNS) [13], and Responsive Neurostimulation (RNS) [14].

VNS has been shown to induce a 31% median reduction in seizure frequency initially, with a long-term 50% seizure frequency reduction rate of 60%, and an 8% seizure freedom rate [15]. Another electrical stimulation alternative for epilepsy is the Anterior Nuclei of the Thalamus (ANT) -DBS which targets the Papez circuit, exhibits a 44% median reduction in seizure frequency initially, with a long-term 50% responder rate of 74%, and an 18% seizure freedom rate [16]. Long-term therapy at 7 years resulted in median 75% seizure reduction [17] RNS implantation and responsive stimulation therapy results in a 42% median reduction in seizure frequency initially, a long-term 50% responder rate of 73%, and a 28% seizure freedom rate [18], and 82% median seizure reduction after only 3 or more years in real-world practice [19]. Despite the demonstrated partial efficacy of these techniques, there exists a substantial margin for enhancing the effectiveness of electrical stimulation protocols in epilepsy management. Development of novel strategies will help improve efficacy of electrical stimulation.

Coordinated Reset (CR) stimulation was computationally developed and aims at inducing long-term desynchronization of abnormal neuronal synchronization by leveraging plasticity mechanisms [20–23]. Neuronal synchronization is relevant under normal as well as abnormal conditions [24]. A number of neurological disorders are associated with abnormal neuronal synchrony, for example Parkinson’s disease [25], tinnitus [26] and epilepsy [27]. Similarly, epileptic seizures are the clinical manifestation of abnormal, excessive, hypersynchronous activity of a population of cortical neurons [28].

The principle of CR stimulation is deeply rooted in the understanding of network dynamics and connectivity. Through a spatiotemporal sequence of stimuli delivered to distinct neuronal populations, CR stimulation aims to disrupt the in-phase synchronization [20]. Initially, CR stimulation was designed in neuronal network models with constant couplings (i.e., synaptic strengths and connectivity patterns) for demand-controlled desynchronization, where CR stimuli were delivered whenever neuronal synchrony reached a certain threshold or CR stimuli were delivered periodically with strength adapted to the amount of neuronal synchrony [20]. The target of the computational approach shifted when spike-timing-dependent plasticity (STDP) [29–31] was incorporated in the computational models [21]. STDP is a basic plasticity mechanism that adapts synaptic weights to the timing relationship of the corresponding neurons, e.g., by increasing synaptic weights if the postsynaptic spike follows the presynaptic one and decreasing synaptic weights in the opposite case [32]. Computationally neuronal networks with STDP may display multistability, where different stable attractor states may coexist [21,33–35]: The neuronal network may remain stable in states with increased synaptic connections and increased neuronal synchrony or in desynchronized states with reduced synaptic weights or in intermediate states with patterned activity patterns, e.g., cluster states. As shown numerically, the tight relationship between network dynamics and connectivity enables to reshape the connectivity by controlling the dynamics [21,36–38]: Desynchronizing stimulation may cause a reduction of coincident firing and, mediated by STDP, a reduction of synaptic weights, in this way shifting the neuronal network from an attractor with strong synchrony and strong synaptic connections to an attractor with weaker synchrony and weaker connections. In this way, the stimulated network may undergo a CR-induced anti-kindling, i.e. unlearning of abnormal connectivity and synchrony [21,37,39,40]. The desynchronizing effects persist after cessation of stimulation. Computationally, it was shown that the anti-kindling principle also applies in the presence of STDP and different types of structural plasticity [41,42]. Computationally revealed qualitative predictions, such as long-lasting and cumulative effects [21,43] were critical for the pre-clinical and clinical development of CR stimulation. By the same token, computational studies predicted that CR stimulation could not only be applied invasively by electrical stimulation, but also non-invasively by different stimulation modalities, e.g., by sensory stimulation [40,44].

Acoustic CR stimulation for the treatment of tinnitus caused a clinically significant reduction of tinnitus symptoms together with a simultaneous decrease of abnormal neuronal synchrony [45,46] and abnormal effective connectivity [47]. A few hours of electrical CR-DBS stimulation led to pronounced weeks-long therapeutic and desynchronizing effects in non-human primates rendered Parkinsonian with MPTP [48–51]. In patients with Parkinson’s disease, electrical CR-DBS delivered to the STN caused a significant and cumulative reduction of abnormal beta band oscillations along with a significant improvement of motor function [52]. Furthermore, non-invasive vibrotactile coordinated reset (vCR) produced no side effects, and delivered sustained cumulative improvement of motor performance [53], which is congruent with the computational findings mentioned above. In a binge-drinking mouse model, electrical CR stimulation of the nucleus accumbens administered during only the initial phase of alcohol exposure or only prior to alcohol exposure significantly reduced binge-like drinking without impairing social behavior or locomotor activity [54].

Relevant to epilepsy, long-lasting desynchronization was induced using electrical CR stimulation in rat hippocampal slices with sustained spiking dynamics using a model of magnesium withdrawal [55]. Electrical stimuli were delivered to different locations of CA3, projecting to different parts of CA1 through the Schaffer collaterals. Local field potentials (LFP) were measured in CA1 with an electrode array. In accordance with computational studies on CR and periodic stimulation [21,56], the experimental study revealed two main effects of CR stimulation [55]: a widespread amplitude decrease in local field potentials (LFPs), reflecting a likely overall decrease in synchrony across several layers of CA1, and a pronounced desynchronization between stratum oriens and stratum pyramidale through distal stratum radiatum. In contrast, periodic control stimulation (with coincident stimulus delivery through all stimulation contacts) induced a long-lasting increase in both overall LFP amplitude and synchrony between stratum oriens and the other CA1 layers.

Building upon this foundational in vitro work, our research study seeks to extend the application of CR stimulation in epilepsy. We used an in vivo animal model of status epilepticus showing sustained epileptiform activity induced through the intrahippocampal application of 4-aminopyridine (4AP) in Wistar rats [57,58]. We used CR stimulation with rapidly varying sequences (RVS-CR), where the stimulation sequence is randomly varied from cycle to cycle [40]. A computational study by Manos et al. (2018)[59] showed that RVS-CR stimulation is more robust against detuning the stimulation frequency relative to the dominant collective frequency component of the network, rendering it more appropriate to epileptiform variations where each recorded signal has a particular spectral signature even under synchronized conditions. Also, computationally it was shown that RVS-CR is more robust in networks with spatially inhomogeneous connections compared to CR with constant or slowly varying sequences [60].

Local Field Potentials (LFP) recordings were acquired from CA1 posterior hippocampus and analyzed to measure acute effects of RVS-CR. LFPs were recorded from several contacts before and after RVS-CR, and differences in spike and oscillatory univariate and bivariate feature metrics were compared to the no-stimulation control condition. Statistical analysis and results showed significant decreases in bivariate spike and oscillatory features across the spectrum for the stimulation group compared to the control condition, in this way motivating further exploration of CR stimulation as a viable intervention for epileptiform activity suppression and hence treatment of DRE.

## 2. Methods

### 2.1 Animal Model of Status Epilepticus (SE)

All *in vivo* animal experiments were approved by the Technion Institutional Animal Ethics Committee. For all animal experiments, adult male wildtype 8-14 week old Wistar rats were used (230-430 gr). Electrographic seizures were induced through local application of 4-aminopyridine (4AP). This chemoconvulsant is a potassium blocker that prevents hyperpolarization, increasing excitability in the brain. 4AP in small doses (0.5 ul with 5 millimolar) can acutely induce recurrent electrographic seizures [57,58,61]. In higher doses (0.5 ul with10 millimolar), 4AP, induces sustained synchronized epileptiform activity resembling Focal and Refractory Status Epilepticus (SE) [62].

During the neurosurgical procedure anesthesia was maintained through inhalation of isoflurane (2.0%) mixed with room air and 100% O2 (1:1 ratio). Animals were positioned on a stereotaxic frame (Model 1900, Kopf, CA, USA). To mitigate basal sensory input during the incision 3 ml of Lidocaine (2%) were injected into the subcutaneous tissue of the scalp. A heating pad maintained the body temperature at 36°C throughout the surgery. A sagittal incision exposed the skull surface, then a craniotomy window, measuring 0.5mm x 0.5mm, was drilled on the right side of the skull overlying the CA1 region of the right posterior hippocampus. Electrode probes were implanted orthogonally to the brain surface utilizing a motorized micromanipulator (MP-285, Sutter Instruments, CA, USA) mounted on a Kopf stereotaxic frame (Model 1900, Kopf, CA, USA). The probes were carefully lowered at a rate of 1mm/60sec, and enabled recording from the entire right posterior hippocampus. To induce seizures and a SE, an intrahippocampal injection of 4AP was administered following an oblique trajectory (65°) into the CA3 region of the right posterior hippocampus (AP -5.8, ML 4.6, DV 5.5) using a glass micropipette attached to a microinjector (Narishige, Japan), held by a motorized micromanipulator (MP-285, Sutter Instruments, CA, USA).

### 2.2 Electrode probes and Histology

Electrode probes were designed in collaboration with NeuroNexus (MI, USA) for this study. The resulting electrode probes attempt to bridge experimental and clinical technology, by resembling depth electrodes used in the clinic. The resultant design, dubbed the rat research Deep Brain Stimulation Array (rDBSA) [63], features four ring contacts dedicated to electrical stimulation alongside ten circular contacts (∅θ = 15 µm) for recording Local Field Potentials (LFP) and Multi Unit Activity (MUA). Recording and stimulation contacts have 500 µm interspacing along 6.5 µm. An illustration of the electrode probe and its implantation site is shown in Figure 1.C.

**Figure 1.**
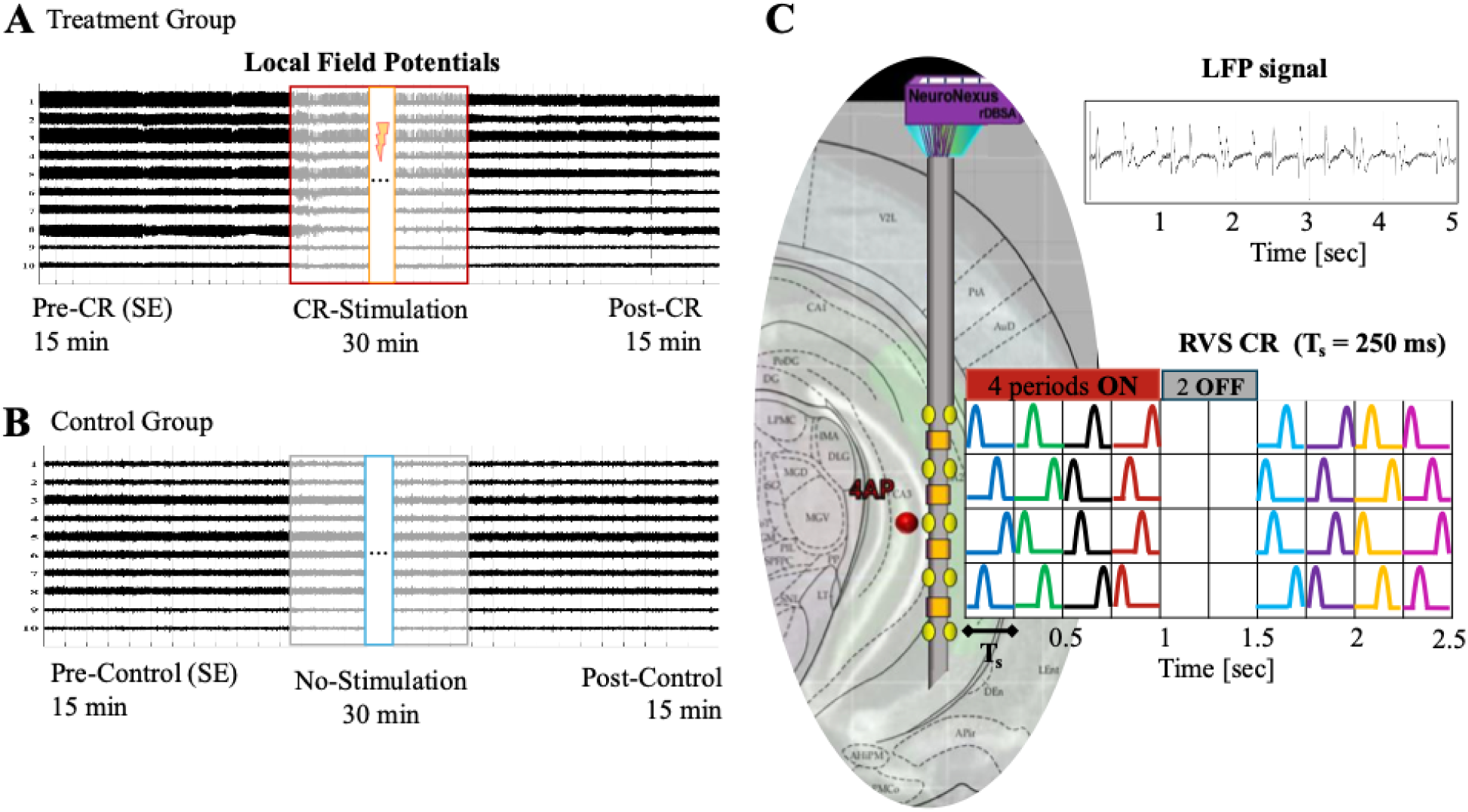
Electrophysiological framework for neural recordings and stimulation. The experimental framework is illustrated for desynchronization of status epilepticus induced by 4AP. In A, the Local Field Potentials (LFP) for ten recording channels before and after CR stimulation is shown. In B, a control recording is shown, where there is no electrical stimulation. In C, we show a graphical representation of the electrode probe in the implantation site at CA1 of hippocampus, with injection site (red disk) of 4AP in CA3. In addition, we show a five second example of the LFP showing sustained epileptiform firing. Next to the electrode probe is an illustration of the Rapid-Varying Sequences Coordinated Reset (RVS-CR) protocol, where the stimulation sites are randomized at each ON stimulation cycle, and four ON periods were followed by two OFF periods. Stimulation period was chosen to be 250 msec.

Following the recording and stimulation procedures, a transcardial perfusion with PBS and 10% paraformaldehyde was performed on the animals to achieve brain fixation. Subsequently, Nissl staining was conducted to verify the accuracy of the electrode placement and, hence, to validate the experimental setup.

### 2.3 Electrographical Recordings and Preprocessing

The acquisition of electrophysiological data was done utilizing a 32-channel AlphaLab SnR system (Alpha Omega, Nazareth, Israel). The system has eight separate stimulation sources alongside user-programmable Digital Signal Processors (DSPs), providing real-time communication for closed-loop applications.

Electrophysiological analog data was filtered for digitalization using a 16-bit ADC in a digitized headstage (Alpha Omega, Nazareth, Israel), which was directly connected to the electrode probe. Data was acquired at a sampling frequency of 11 kHz. Preprocessing of LFP recordings included downsampling signals to 1000 Hz, notch filtering for line-noise at 50 Hz, and bandpass filtering between 0.5 and 100 Hz with a fourth order Butterworth filter. LFPs were referenced to a ground wire on the surface of the cortex. A 5-second recording example is shown in Figure 1.C. Non-neural electrographic artifacts (characterized by signal saturation were excluded from analysis.

Electrographical recordings from eight animal subjects were used for analysis in this study, five animals were subjected to CR stimulation and three were used as stimulation-free controls. Each recording is composed of two epochs of 15 minutes each. For the CR stimulation group, the first epoch is taken from LFP recordings during SE and precedes CR stimulation (pre-CR), and the second epoch begins at the offset of CR stimulation and similarly lasts 15 minutes (post-CR), this is illustrated in Figure 1.A. For the Control group the epochs are recorded similarly, however the electrical stimulation is replaced by a stimulation-free pause, as illustrated in Figure 1.B.

### 2.4 CR Stimulation

Following the 15 min pre-CR baseline epoch of SE recording, electrical CR stimulation was delivered in periods of length Ts through the four ring stimulation contacts in coordinated sequences. CR was administered for 30 min, adhering to a 4:2 on:off CR pattern, where 4 on-stimulation periods were intersected by 2 off-periods, with constant stimulation period Ts [64] (Fig. 1C). We used a 4:2 on:off CR pattern because in a computational study the intermingled off-periods enabled an effective desynchronization while reducing the integral amount of stimulation [64]. In addition, nearly all pre-clinical and clinical CR studies used the intermingled on:off CR pattern [45–55]. During each on-stimulation period a sequence of 4 electrical stimuli was delivered, where each stimulation site was activated exactly once. Single stimuli were delivered in equidistant steps of Ts/N, where N = 4 denotes the number of ring contacts used for stimulus administration (Fig. 1C). A computational study on optimal number of stimulation contacts confirmed that 4 stimulation sites are sufficient for effective CR neuromodulation [64]. The single stimuli were composed of a pulse-train of six charge-balanced pulses with an intra-burst frequency of 120 Hz, pulse amplitude of 200 *µ*A, and a pulse width of 100 *µ*sec. The CR stimulation frequency f = 1/Ts was chosen to be close to the spike firing frequency during SE which in the pre-CR baseline was close to 4 Hz (4.05±0.45 Hz) for all subjects. Accordingly, we selected f = 4 Hz in all subjects. We administered CR stimulation with rapidly varying sequences, where the sequence order of contact sites used for stimulation was randomized with equal distribution from one period to another with equal probability (Fig. 1C). RVS-CR stimulation was used due to its computationally demonstrated robustness against variations in the stimulation frequency (relative to the collective network frequency), its effectiveness when stimulation frequencies are in the range of neuronal firing rates and its robustness in the presence of spatially inhomogeneous connectivity patterns [59,60].

### 2.5 Electrophysiological Data Analysis

A sliding window analysis was performed on the preprocessed data, using two-second non-overlapping windows to extract features sensitive to neural synchronization for analysis of recording periods. Time-series features included the spike rate, spike and continuous signal amplitude and bivariate spike relative phase. Time-frequency representation features of the continuous signal included univariate power and bivariate phase-difference between channel pairs in the following frequency bands:

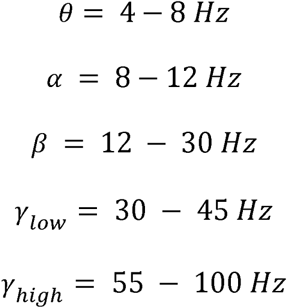

#### 2.5.1 Spike Event Analysis

##### Spike amplitudes

Spikes were identified using the derivative of the signal by identifying the peak at the change in sign. To avoid undesired detections a minimum threshold value to cross was imposed. This threshold value was set adaptively for each window, either being the 95 percentile of the analysis window or for the entire recording, whichever one was greater. Signal amplitude was computed as the maximum value of the signal minus the minimum value of the signal for that time window. Spike peak amplitude distributions pre-CR and post-CR were separately analyzed by channel to detect spike modulation. An example of the amplitude analysis done for all channels is shown in Supplemental Figure 1.

##### Relative spike phase

A bivariate relative phase analysis was also performed on the spike series for each channel. In this analysis the spike events (spike peaks) in channel *m* (*m = 1,..,10*) are observed at times t_m,i_, where *i,* denotes the spike event in the time-series. For each spike-period, i.e. the period between spike *i-1* and spike *i* for channel *m,* we identify all spike-events in the remaining channels that fall within that spike-period. The time difference between spike-events is calculated for each spike-period and then the spike relative phase for each channel pair is calculated in the following manner:

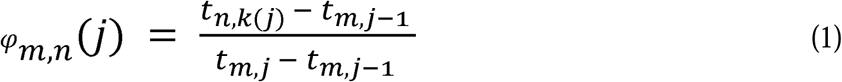

Where φ*_m,n_ (J)* has a value between 0 and 1 and can be used to detect modulation of the spike relative phase between channel pairs (*m, n*). For analysis, we constructed matrices in which rows represented the reference channel (*m*) and calculated the relative phase to all other channels, n, as columns. These matrices contain information on how spike events (j) across channel pairs are mutually related using several references to identify coordination in network subpopulations. This is used to analyze CR stimulation-induced modulation in spike-events between the pre-CR to post-CR periods. An case of the relative spike phase analysis is shown for subject CR-stim 1 in Supplemental Figure 3.

#### 2.5.2 Spectral Analysis

*Spectral power:* The spectral power for the five different frequency bands was calculated using Welch’s method, with Hamming windowing [65]. The spectral heatmap of the LFP is shown in Figure 2.B for one of the experiments. For each of the 10 channels, the mean power for each frequency band was computed for both pre-stim and post-stim periods. Then the percentage change in power for each frequency band was calculated using the following formula 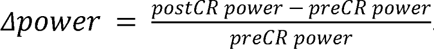. In Figure 2.C, the blue bars represent the power for the pre-CR period and the red bars the power for the post-CR period for each frequency band. Figure 2.D shows the result of the average for each frequency band for all channels, this was then repeated for all animal subjects (Supplemental Figure 2.). For univariate group analysis, the percentage changes were then averaged across channels and then compared between control and CR stimulation group using permutation tests.

**Figure 2.**
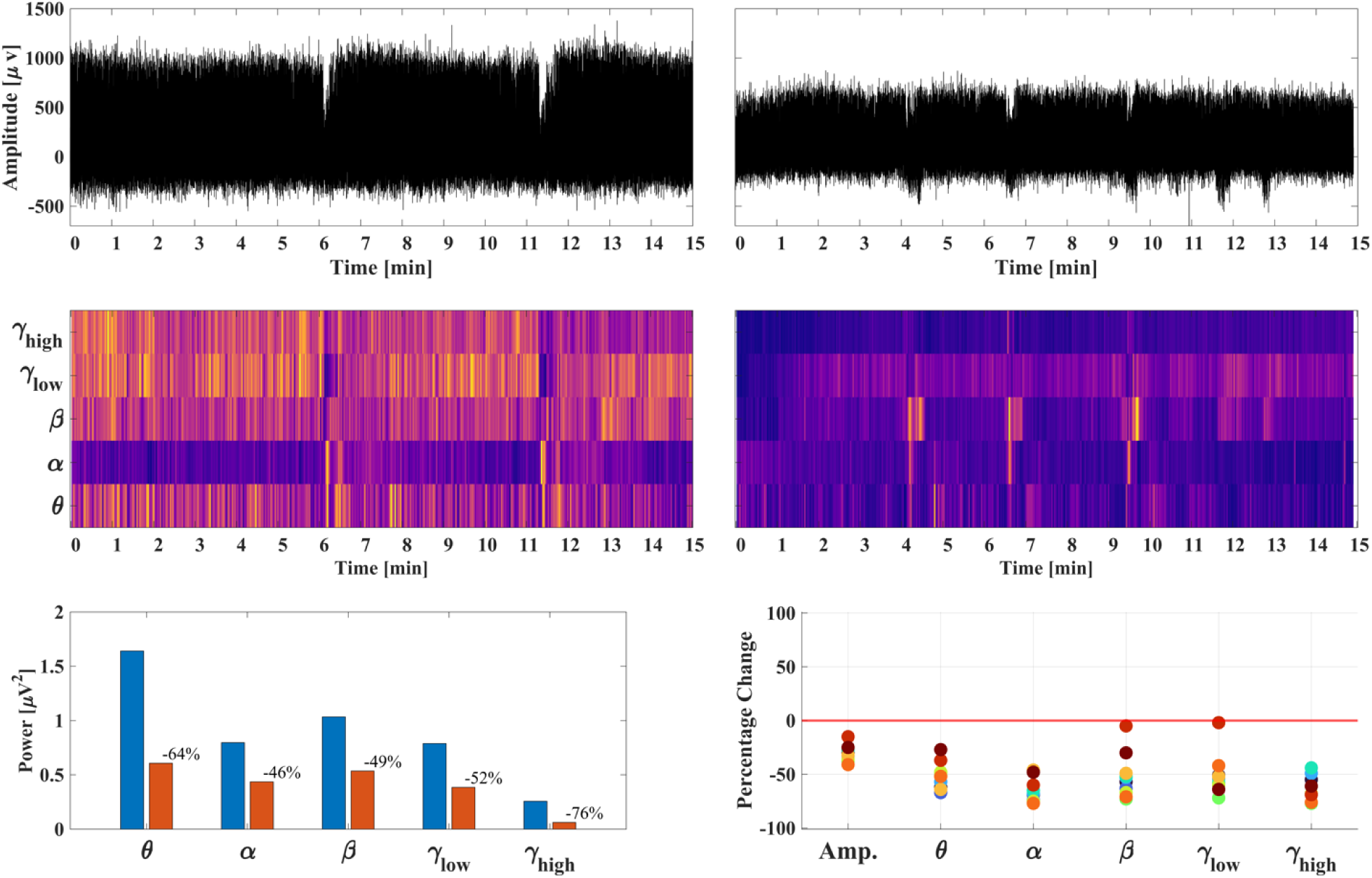
Univariate feature analysis. The processing pipeline for univariate features is illustrated. In A the pre-CR and post-CR electrographical recordings are shown for a single channel. In B, the time-frequency representation of the LFP is illustrated in a spectral heatmap for all frequency bands. Then the percentage change for each univariate feature is calculated. This is shown in C, the blue bars represent the power for the pre-CR period and the red bars the power for the post-CR period for each frequency band. This process is done for each channel. D shows the result of the average for each frequency band for all channels.

##### Time-continuous Hilbert phase

The stroboscopic phase defined in Equation [1] was used to characterize synchronization processes involving the compound spike signal. In addition, we also determined continuous phases of the different frequency bands separately by means of the Hilbert transform [66]. In general, the presence of phase synchronization between two continuous signals within a time window is characterized by the distribution of their phase difference values within this time window [67,68]. While the Phase Locking Value (PLV)[69] is commonly used to measure synchronization, it relies on the circular mean of the phase difference distribution, which can produce spurious results for non-unimodal distributions. Since we observed that not all phase distributions were unimodal, we instead used the general definition of phase synchronization, characterized by one or more pronounced peaks in the circular phase distribution [67,68]. For the bivariate phase analysis, we first calculated the corresponding time-continuous phases of the LFP signals for all frequency bands by bandpass filtering and then computing the Hilbert transform. Second, we calculate the phase differences for all LFP pairs with the different frequency bands, respectively, for pre-CR and post-CR periods separately. The phase difference distribution analysis for subject CR-stim 5, beta band are shown in Supplemental Figure 5 to illustrate that not all phase distributions were unimodal.

#### 2.5.3 Functional Connectivity Analysis

A Functional Connectivity (FC) matrix is created using the bivariate features obtained by the Kuiper statistic. In order to quantify significant differences or coordination for each channel pair the Kuiper circular statistical test was employed [70]. The Kuiper test was shown to be useful, e.g., in Tass et al. 2004, Baud et al. 2018 [71,72] as a circular statistical test, that is computationally efficient and yields a statistic value between 0 and 1, making it practical for intersubject comparative analysis. In particular, the Kuiper statistic does not require the distribution of the phase difference to be unimodal. We calculated the Kuiper statistic for all channel pairs to construct a FC matrix of phase coordination. The Kuiper test [70] uses the cumulative phase distribution and was used on the phase difference between channel pairs. The 1-sample test was used to test against uniform circular distribution and follows the following formula

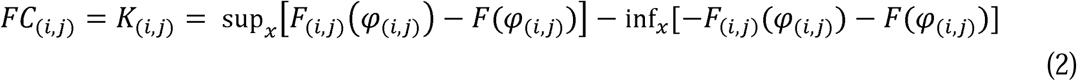

Where the Kuiper statistic for channel pair *i,j* is calculated using the cumulative distribution function of the phase difference φ(φ_(*i,j*)_), which is equal to 1 if <_(*i,j*)_ <x and equal to 0 otherwise.

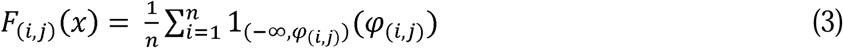

The 2-sample Kuiper test was used to compare the pre-post-CR circular distributions on the spike relative phase and the time-continuous phase difference distributions for each frequency band. In this case the 2-sample test was used to detect pre-post-CR changes of the stroboscopic as well as time-continuous phase differences. In addition, to detect synchronization patterns in the pre-CR and post-CR periods separately, the 1-sample Kuiper test (comparison with uniform circular distribution signifying complete absence of phase synchronization) was calculated for the phase distributions of each channel pair and each frequency band for each period (i.e. pre-CR and post-CR). The FC for the spike relative phase using the 2-sample Kuiper test is shown in Figure 3. The 1-sample and 2-sample Kuiper FC matrices for the spike relative phase and phase synchronization in all frequency bands are shown in Supplemental Figures 4, 6 and 7.

**Figure 3.**
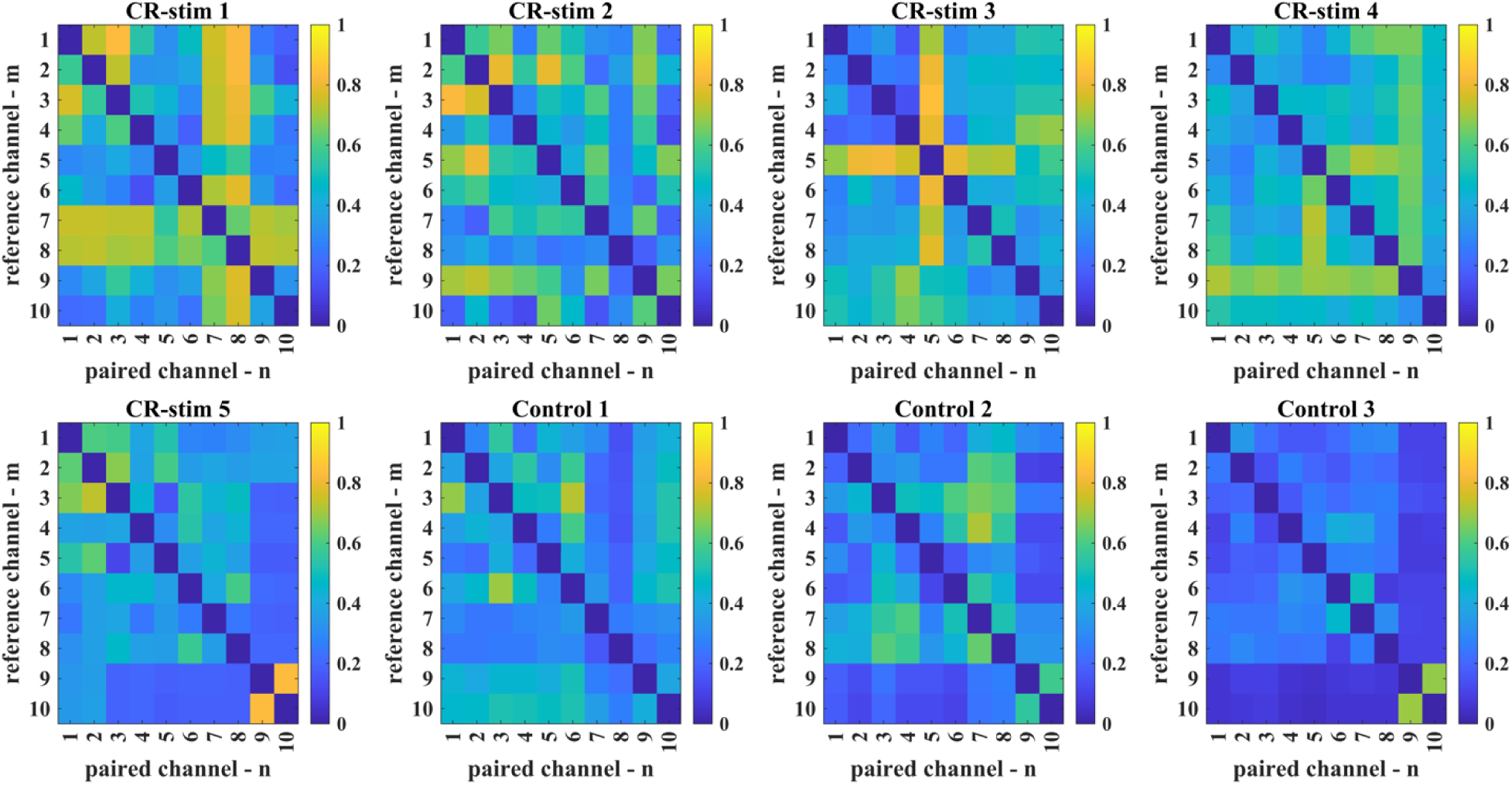
Functional connectivity matrices: 2-sample Kuiper test. Functional Connectivity (FC) matrices are constructed for all subjects, CR stimulation (N=5) and control subjects (N=3). The Spike relative phase feature matrices are derived from the electrographical recordings pre and post periods for each subject. The 2-sample Kuiper test is then used on the pre and post feature matrices to construct a single 2-sample FC matrix for each subject.

#### 2.5.4 Network Connectivity Analysis

To analyze network changes based on functional connectivity, we computed graph theoretic network measures of functional segregation and integration as proposed in [73]. These network measures offer critical insights into the structural and functional organization of brain networks, revealing patterns of neural communication and integration. Their analysis provides insights into both the organization, efficiency adaptability and the dynamics of neural systems [74]. In this study the network measures were computed directly from the FC matrices (2-sample Kuiper statistic, 1-sample Kuiper statistic), as they are a weighted non-directional connectivity matrix. Binary connectivity matrices were also derived from the FC matrices by thresholding the Kuiper statistic value, which served as the adjacency matrices for the analysis. A threshold sweep was done to calculate the binary connectivity matrices – adjacency matrices from 0.1 to 0.9 in steps of 0.05. This was done to identify specific ranges of network connectivity measures that might be more descriptive of perturbation effects, this also provides insight into the resolution limit of the network measure. Network measures of functional segregation and integration were derived from the FC matrices and used for analysis. To evaluate network desynchronization, i.e., desynchronization between different recording sites, we focused mainly on network segregation measures. The measures used to evaluate functional segregation were:

- Average Degree of the Network: The average degree reflects the number of connections each node has, averaged across all nodes. It quantifies the network’s overall connectivity density and indicates the potential for communication between regions [73].
- Average Clustering Coefficient: This measure quantifies the degree to which nodes in a network tend to cluster together. It is calculated as the fraction of a node’s neighbors that are also neighbors of each other, indicating local connectivity [75].
- Transitivity: A global variant of the clustering coefficient, transitivity measures the proportion of all possible triangles in a network that are actually present. It provides an indication of how well connected the local neighborhoods of nodes are on a global scale [76].
- Modularity: Modularity measures the degree to which a network can be divided into clearly defined, non-overlapping sub-networks or modules. High modularity indicates the presence of dense connections within modules and sparse connections between them, which is a hallmark of functional segregation [77].

Since CR stimulation aims at desynchronizing the network, these functional segregation quantities measure the changes in clustering tendencies between the nodes, based on the FC matrix, describing the functional connectivity effects of CR stimulation. To track the efficiency of information flow in the network we also calculated network measures evaluating functional integration:

- Betweenness Centrality: This measure quantifies the importance of a node in terms of the number of shortest paths that pass through it. Nodes with high betweenness centrality serve as key intermediaries in communication and are often critical for efficient information flow across the network [78].
- Average Shortest Path Length: This is the average number of steps along the shortest paths for all possible pairs of nodes. It represents the efficiency of information flow within a network. Shorter average path lengths suggest faster and more efficient communication between regions [75].

Functional integration measures describe how different regions of the brain coordinate and communicate, reflecting how efficiently information is exchanged across the whole network. In addition to these measures, we computed the Frobenius norm and maximum eigenvalue of the weighted FC matrices as a general metric of overall network synchronization. In Figure 4, the 2-sample spike relative phase FC matrix is shown as an electrode network graph depicting a weighted undirected connectivity matrix. Figure 5 illustrates the corresponding percentage change calculation done on the 1-sample Kuiper FC matrices.

**Figure 4.**
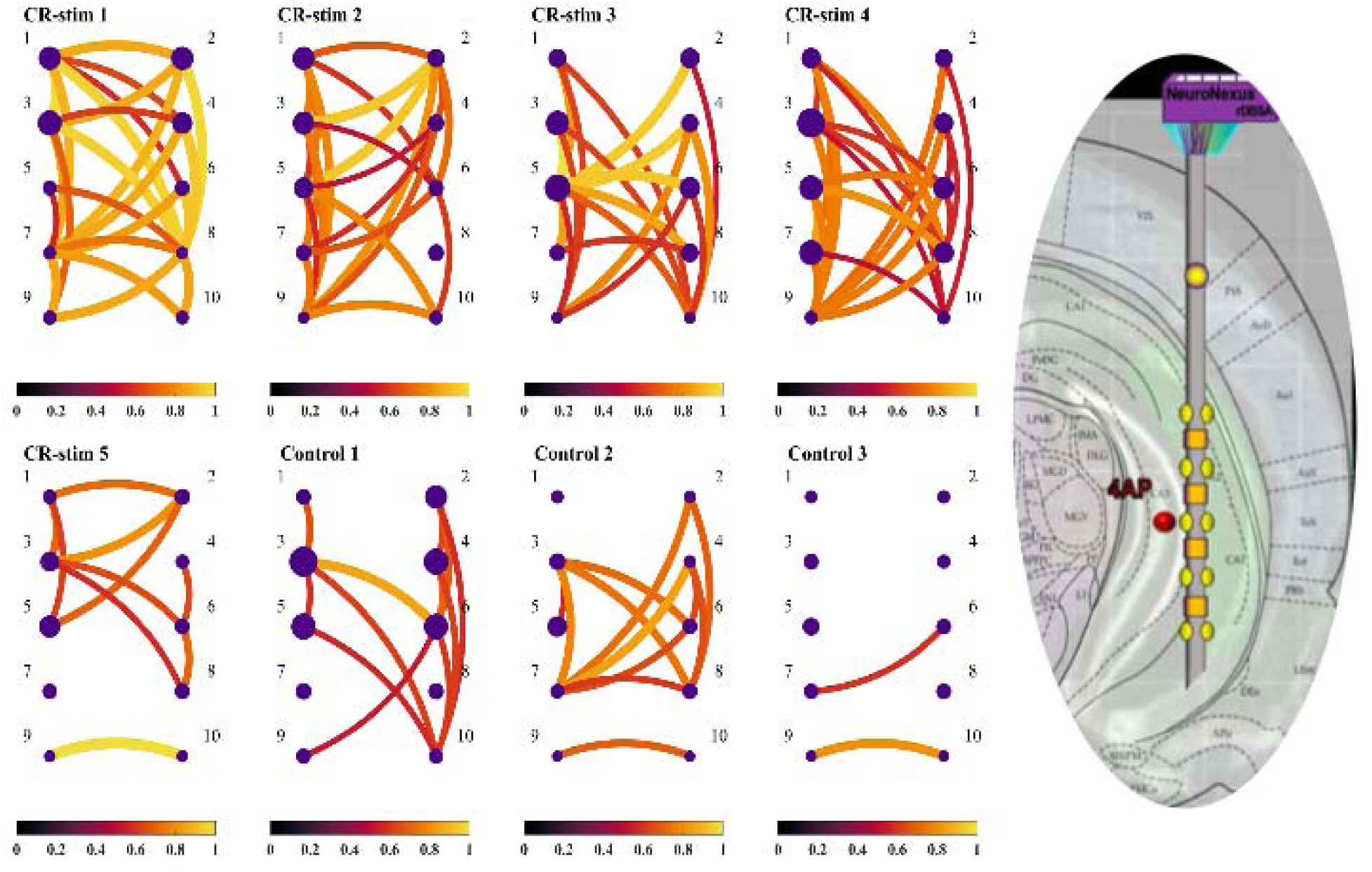
Electrode network connectivity graph. The 2-sample weighted functional connectivity matrices are used to build network connectivity graphs for all features, in this case for the spike relative phase. To the far right the illustration of implanted electrode is shown as an inspiration for creating these weighted undirected network graphs. Graphs are constructed for all subjects, belonging to CR stimulation (N=5) and control group N=3). The diameter of each node represents the average electrode continuous signal amplitude. The color and width of each edge weight is representative of the Kuiper statistic and strength of connection.

**Figure 5.**
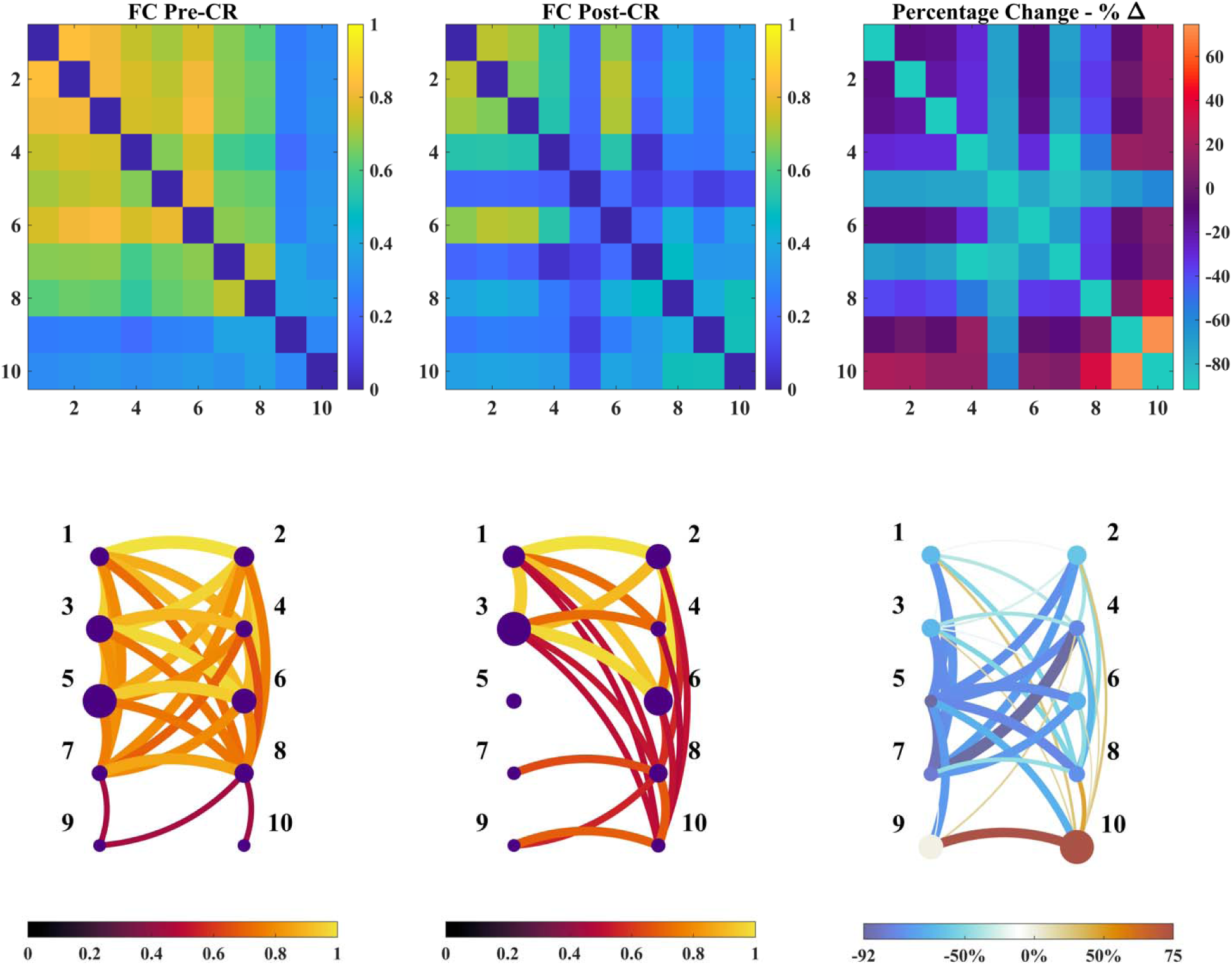
Functional connectivity state differential matrices and connectivity graphs. The 1-sample weighted functional connectivity matrices and their percentage change is shown for subject CR-stim 3, alpha band. On the top row are used to build network connectivity graphs as seen in the bottom row. The diameter of each node represents the average electrode continuous signal amplitude. The color and width of each edge weight is representative of the Kuiper statistic and strength of connection. For the percentage change connectivity graph, blue marks a decrease and red represents an increase. Blue nodes show channels that had a decrease in signal amplitude, blue edge weights represent a decrease in FC for that channel pair, and in red edge weights that increased.

### 2.6 Statistical Analysis

To test for differences between the CR stimulation and control groups, permutation tests calculating the effect size and Cohen’s D were used computed and empirical p values calculated. Employed tests involved 10,000 permutations for each condition and each feature measure. These series of permutation tests were employed to establish potential significance between the CR stimulation and control group in this exploratory study. Empirical p-values <0.05 were considered statistically significant. Because the exploratory nature of this study, p-value correction for multiple testing was not performed; the goal of this study was hypothesis generation rather than formal testing.

The permutation tests as described above were done on the 2-sample FC matrices derived network measures (e.g. FC norm, clustering coefficient). To analyze the 1-sample FC matrices network measures, we applied two statistical approaches: (i) Computing the relative percentage change in network measures from the pre-CR to the post-CR period. Then the permutation test was done between the CR stimulation and control group percentage change to test significant differences between groups. (ii) A two-way factorial analysis (TWFA) between the pre-CR and the post-CR period for the CR stimulation and control groups. In (ii) we tested for the treatment effect, the time effect and the interaction effect, using the pre-CR and post-CR periods as well as on the pre-no-stim and post-no-stim periods. The treatment effect was calculated by the difference of the sum of all periods between CR stimulation and control group. The calculation of the time effect was done by the difference between pre-CR and post-CR periods. Finally, the interaction effect was calculated as the difference in the mean change between pre- and post-CR periods for the CR stimulation group, compared to the same mean change for the control group. The two-way factorial effect calculation and significance assessment was done using permutation tests for each effect as described above, where the Cohen’s D was calculated over 10,000 permutations. Results for all permutation tests are shown in Supplementary Tables 1-4. Then a graphical summary and visualization for all significant permutation test results is shown in Figure 6. All computational analysis scripts were custom written in Matlab (Mathworks, Naswick, MI, USA), Network measures were calculated using the Brain Connectivity Toolbox [79], code for signal acquisition and stimulation was written using the Alpha Omega API.

**Figure 6.**
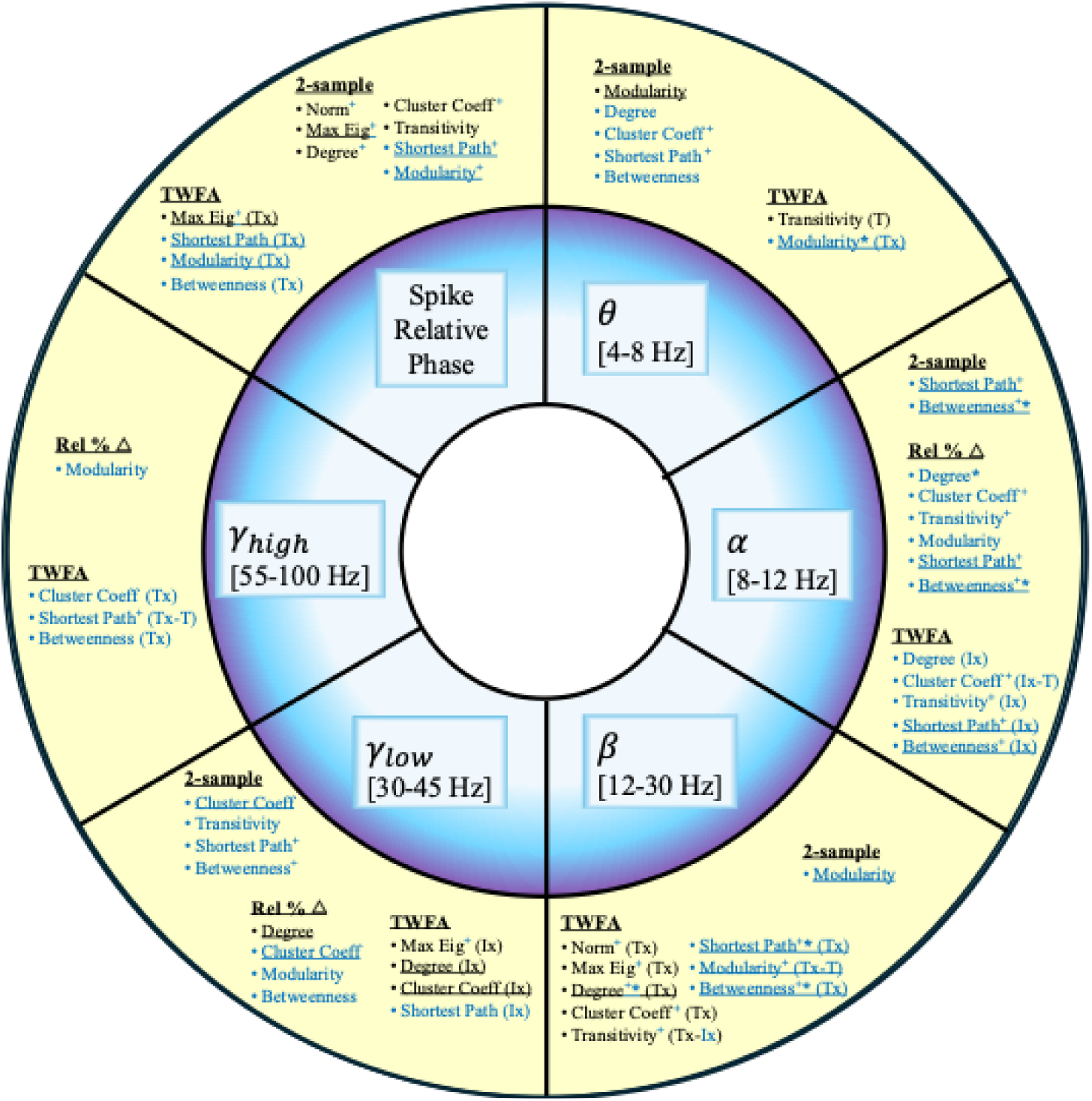
Comprehensive permutation test analyses results. This figure highlights for each frequency band and spike relative phase, which feature metrics were found to be significant using different statistical analyses between the CR stimulation and the control group. The 2-sample feature metrics, and 1-sample relative percentage change and Two-Way Factorial Analysis significant feature metrics are listed accordingly. In **black** metrics that were significant with alpha = 0.05 for the weighted FC matrix. In **blue** metrics that were significant with alpha = 0.05 for a thresholded binary FC matrix. All metrics in black also had a significant binary FC matrix. Metrics that were significant in two statistical analyses are <u>underlined</u>. **+** Indicates that the feature metric was significant for more than one threshold. **\*** denotes significance with alpha = 0.01. Two-Way Factorial Analysis (TWFA) effects are abbreviated as such: Time effect – T, Treatment effect – Tx, Interaction effect – Ix.

## 3. Results

We analyzed the 15 min pre- and post-CR epochs that surrounded the 30 min CR stimulation period. Both pre-CR and post-CR epochs were quantified with different univariate and bivariate measures (see Methods), and their differential analyzed between the CR stimulation group and the control group. Quantitative analysis of the pre-CR and post-CR periods involved the extraction of univariate features from single channels and bivariate features computed from channel pairs. The univariate features included the spike amplitude (of the compound LFP signal) as well as the continuous signal amplitude and the spectral power for different frequency bands. The bivariate features extracted included the relative phase of the spikes as well as the phase difference between all channel pairs within all frequency bands. From the phase difference a circular statistic was computed by means of the Kuiper statistic in two different ways: (i) We detected significant patterns of phase synchronization in each (pre- and post-CR) period separately, using the 1-sample version of the Kuiper test for pre-CR and post-CR periods separately: Comparison to a uniform circular distribution enabled to reveal phase synchronization patterns in each period separately. (ii) We used the 2-sample Kuiper test to detect changes of the amount of phase synchronization between pre- and post-CR epochs. The circular statistics were used to detect significant changes between the phase difference distributions between the pre-CR and post-CR epochs. These circular statistics derived from bivariate features were then used to construct FC matrices. Functional connectivity was analyzed by computing network connectivity measures and using statistical permutation tests to assess significant modulation between pre-post periods for CR stimulation and control groups.

Each feature was used to compare modulation between the pre-CR and post-CR epochs for each animal subject. Feature measures were then compared between the CR stimulation and control groups to determine the efficacy of CR stimulation in modulating sustained epileptiform activity.

The univariate spike analysis entailed calculating the spike rate and amplitude for all channels for each epoch and then averaged across channels and calculated the pre-vs. post-CR epoch differential. Univariate features for the pre-CR and post-CR epochs were obtained and their differential percentage changes were calculated as shown in Figure 2. This process was repeated for all channels and all animal subjects and, ultimately, grouped according to CR stimulation or control conditions.

### Mean spike rate

The mean pre-CR spike-rate for all recording contacts and all subjects was 4.045+0.44 (mean+std), and for the post-CR epoch a spike rate of 3.93±0.45 was measured. There was no significant difference between pre-post epochs or between subject groups.

### Amplitudes

Amplitude results showed a marked decrease across channels for all subjects in both groups. This was measured for the continuous signal amplitude and the spike amplitude. The spike amplitude distributions by channel were compared between the pre-CR and post-CR epochs. The spike and continuous amplitude distributions by channel were compared between the pre- and post-periods, by computing the average percentage change between post-pre periods. Supplemental Fig 1 shows an example of the distribution of the spike amplitudes for all channels for a subject. For the continuous signal amplitude, the CR-stimulation group had an average decrease of -23.4±7.46%, and the control group presented a decrease of -11.7±7.46%. Similarly, the spike signal amplitude for the CR-stimulation group showed a decrease of - 25.3±7.34%, and the control group -14.5±7.71%. Regardless of the larger decrease in amplitude in the CR-stimulation group, neither of the amplitude metrics had a significant result using the permutation tests comparing treatment and control groups.

### Spectral power

Time-frequency-representation of the LFP using Welch’s method was done to calculate the power in 5 frequency bands (*θ, α, βγ, γ_low_, γ_high_*). For illustration, a resulting spectrogram is shown in Figure 2.B. Univariate results from the power differential percentage are shown in Supplemental Figure 2, for all subjects we observed a marked decrease in spectral power for all frequency bands in most channels. For control subjects there was an overall decrease in features but less marked than with the CR stimulation group. Statistical analysis with permutation tests did not reveal any significance for the average percentage change in univariate power measures between CR stimulation and control groups.

### Relative spike phase

We assess compound synchronization patterns of the 10 LFP channels based on the pairwise phase relationships of their spikes. To this end, first, we considered whether there are any relevant changes of the synchronization patterns between pre-CR vs. post-CR epoch (for the CR stim group) as well as between before and after the 30 min no-stim epoch (for the no-stim control group). We performed a bivariate relative spike phase analysis using the 2-sample Kuiper statistics for all LFP channel pairs, comparing pre and post relative spike phase distributions, respectively. This revealed that there are different levels of modulation across animal subjects. This is illustrated in Figure 3, where there is an observable difference in Kuiper statistic values between the CR stimulation subjects and the no-stim control group. The CR stimulation subjects show higher value than the control group for all feature metrics derived from the spike relative phase.

### Phase synchronization within frequency bands

We applied the same two-step synchronization analysis strategy to all LFP channel pairs in each frequency band. First, we compared the amount to which the compound LFP pairs’ synchronization patterns changed in the CR-stim vs the no-stim group. Second, we assessed the direction of these change, e.g., overall increase or decrease of synchronization. To this end, in analogy to the relative spike phase analysis, we applied the Kuiper statistic for the analysis of the change of the phase difference distribution when comparing pre-vs post CR and pre-vs post no-stim pause for the different frequency bands (Supplemental Figures 6, 7).

Using the 1-sample Kuiper test was performed on the pre-CR and post-CR conditions as well as on the pre-no-stim and post-no-stim conditions separately. In this way, for the four different conditions (pre-CR, post-CR, pre-no-stim, post-no-stim) we detected phase synchronization by testing the observed phase difference distribution against a uniform circular distribution, representing complete desynchronization [Tass et al. 1998, Rosenblum et al. 2001]. Finally, the 2-sample Kuiper test on the pre-CR and post-CR as well as the pre-no-stim and post-no-stim channel-pair phase differences for all frequency bands, each tests’ statistic values were grouped and used as a modulation metric. These findings show that there is a significant change in the phase distribution of different frequency bands for the CR-stim group between the pre-CR and post-CR conditions compared to the control group. The CR stimulation subjects showed a lower value than the control group for all feature metrics derived for continuous signal phase synchronization for all frequency bands.

### Functional network connectivity

The bivariate Kuiper test matrices were employed to evaluate phase coordination across different frequency bands, constructing FC matrices from both spike and continuous signals. The FC matrix for each frequency band was generated by comparing pre- and post-stimulation periods, using the 1-sample Kuiper test for intra-period phase coordination and the 2-sample Kuiper test to quantify changes across periods (Figures 4,5). The output of these tests represents a weighted connectivity matrix where each element reflects the strength of phase coordination. From these weighted FC matrices, binary networks were constructed through thresholding, followed by sweeping over a range of threshold values to capture meaningful network properties across different resolution limits. Following the construction of FC matrices, complex network measures were computed to quantify functional segregation and integration, following the methods outlined in Rubinov and Sporns (2009)[73]. Metrics such as network degree, clustering coefficient, transitivity, and modularity provided insights into network segregation, while measures like average shortest path length and betweenness centrality described integration.

Statistical permutation tests: To ensure robustness, we applied permutation tests with 10,000 iterations to compare CR-stimulated and control groups. We calculated the Cohen’s D, a measure of effect size, to assess statistical significance (Supplementary Tables 1-4). Permutation tests were done on the 2-sample FC matrices, and for the 1-sample FC matrices we performed the permutation test on the relative percentage change (i.e. pre-post conditions). Permutation tests were also employed in a TWFA using the four different conditions (pre-CR, post-CR, pre-no-stim, post-no-stim). These tests were done for the weighted connectivity matrices and for the binary threshold sweep connectivity matrices. This exploratory statistical framework was designed to capture changes and modulation in network dynamics and desynchronization induced by CR stimulation.

Assessment of effect sizes based on Cohen’s D revealed encouragingly large values for a number of statistical tests (see Supplemental Tables 1-4). All significant results are condensed in Figure 6 for illustration. For bivariate phase-based measures using a weighted FC matrix, significant findings were detected for the relative spike phase. Two-sample tests revealed significance for the FC norm, maximum eigenvalue, and network metrics such as average degree, clustering coefficient, and transitivity. Transitivity increased in the control group but decreased in the treatment group. This suggests greater network desynchronization and reduced clustering in the CR-stimulated group. The decreases in FC norm and maximum eigenvalue reflect a loss of phase coordination, indicating more substantial modulation of spike synchronization in the treatment group. The theta frequency band showed significant increases in modularity for the treatment group and decreases in the control group. Transitivity was also significant between groups although it decreased for both, likely reflecting 4AP washout effects, which leads to more synchronized neuronal activity.

The alpha frequency band showed statistical significance for the binary connectivity matrices, showing high significance between CR stimulation and control for functional integration network metrics (i.e. shortest path and betweenness centrality) for the three statistical analyses.

In the beta frequency band, significant reductions in FC norm, maximum eigenvalue, and network metrics were observed in the treatment group, while the control group showed an increase, except for FC norm and transitivity, which also decreased. These results indicate that CR stimulation reduced network synchronization and connectivity in this frequency band.

For the lower gamma frequency band, interaction effects showed significant decreases in the maximum eigenvalue, average degree, and clustering coefficient for the treatment group, while the control group exhibited increases. This suggests that CR stimulation disrupted phase coordination and reduced epileptiform network connectivity in this band.

Finally, the threshold sweep analysis of binary FC matrices confirmed significant changes across all frequency bands, especially in the delta, alpha, beta, and lower gamma bands. Significant network measures for this frequency bands consistently included both functional integration (i.e. average shortest path and betweenness centrality) and segregation metrics (i.e. average degree and clustering coefficient), among others (See blue metrics in Figure 6). The alpha and lower gamma band showed more significance using the relative percentage change permutation tests, while the other bands showed more significance using the 2-sample (i.e. theta) and the TWFA (i.e. beta). Although both groups showed decreases in these metrics between pre-post periods, the CR stimulation group showed a significantly larger decrease in functional segregation metrics generally, signifying greater desynchronization validated through several features and statistical tests. Conversely, the CR stimulation group showed higher values for significant functional integration metrics in the alpha, beta and lower gamma frequency bands. The observed desynchronization across multiple frequency bands highlights the effectiveness of CR stimulation in modulating neural synchronization. These results strongly support the potential of CR stimulation in attenuating pathological network synchronization, particularly in the beta and lower gamma frequency bands.

## 4. Discussion

The presented findings shed light and continue to build upon previous epilepsy studies (Tass et al. 2009), supporting the promising potential of Coordinated Reset (CR) stimulation in modulating neural synchronization of sustained epileptiform activity and as a treatment for DRE. Main univariate analysis results showed an overall, yet not statistically significant decrease in signal amplitude and spectral power across all frequency bands. Particularly, significant modulation was found in bivariate feature measures for all frequency bands in the CR-stim group compared to the control group, as summarized in Figure 6.

The substantial and significant decrease in spike relative phase and network connectivity in the CR stimulation group in contrast to control conditions, might suggest neuromodulation and desynchronization of the status epilepticus. Additionally, the significant difference in FC matrix norm and max eigenvalue, for the beta and lower gamma frequency bands extends the possible efficacy of CR stimulation in modulating neural synchronization. This observation hints at a broader spectrum of frequency bands that could be influenced through CR stimulation, thereby suggesting its versatility in clinical settings.

The consistency in spike rate pre and post CR stimulation reflects a reliable and reproducible experimental model. This speaks to the rigor of the developed Status Epilepticus animal model and the CR stimulation protocol applied, strengthening the methodological foundation of the study. The utilization of the Rat Research Deep Brain Stimulation Array (rDBSA) and the high-resolution AlphaLab SnR system enriches the methodological rigor, providing a nuanced understanding and precise manipulation of neural dynamics, which is crucial in bridging the experimental to clinical application gap.

Circular statistics were used to analyze channel pairs in a bivariate manner. Specifically, the Kuiper test was used as it has been shown to be appropriate for the analysis of circular data [71,72,80]. The bivariate analysis of the spike relative phase for the pre-post CR periods, detected stimulation-induced changes of relative phase between the spike events of channel pairs. Additionally, to test phase coordination in each period separately, we used the circular statistics in 1-sample tests against a uniform circular distribution. This was done on bivariate features from the phase difference between channel pairs for all frequency bands. To test phase difference modulation between the pre-CR and post-CR periods we used the 2-sample Kuiper test. Since it is a 2-sample test between the pre-CR and post-CR epochs, then higher test statistic values mean a more pronounced modulation between periods. This would mean that the phase difference distribution between channel pairs for all frequency bands shows greater modulation in the CR stimulation group than the control group. This can be understood as desynchronizing or disrupting ongoing patterns and functional couplings that arise in the presence of sustained epileptiform activity.

To analyze the effects and modulation induced by CR stimulation on these functional couplings, we computed network connectivity measures to quantify network changes of functional segregation and integration [73,74]. Functional segregation measures the degree to which brain regions form specialized, locally connected clusters or modules that process specific types of information independently of other regions. Functional integration measures the extent of coordinated communication between different brain regions, reflecting how efficiently information is exchanged across the whole network [73]. These measures have significantly advanced our understanding of brain networks, particularly in neurological conditions like epilepsy, schizophrenia, autism, Alzheimer etc. [74,81]. By examining how these networks become imbalanced, we can better understand disruptions in brain function. Specifically, interventions such as electrical stimulation can be used to modulate the balance between functional integration (coordination across brain regions) and segregation (specialized processing within regions). In epilepsy, restoring and preserving this balance is crucial to prevent seizures [82–85], in particular, in focal status epilepticus where periodic discharges do not resolve with conventional antiseizure medications and anesthetics [86].

Our results showed a general decrease in feature measures, although the reductions in spike and continuous signal amplitudes were not statistically significant in permutation tests, this reflects the robustness of our model and rigor of our statistical framework. Similarly, decreases in average univariate power across frequency bands were observed but lacked statistical significance.

In terms of functional connectivity (FC), significant findings were detected for the bivariate measure of relative spike phase. With the CR stimulation subjects showing higher value than the control group for all feature metrics derived from the spike relative phase. This indicates that, in general, for pairs of channels, the measured relative spike phase between identified spikes consistently exhibited greater changes across all network features, implying spatial-temporal spike decoupling across the network. Specifically, two-sample tests revealed significance for FC norm, maximum eigenvalue, and network metrics including average degree, clustering coefficient, and transitivity. Transitivity, however, increased in the control group. These findings indicate a significant reduction in network clustering and connectivity in the treatment group compared to the control group. The decline in spike relative phase norm and maximum eigenvalue suggests a loss of spike coordination within the network, pointing to greater modulation of spike phase synchronization in the CR-stim group compared to the control group.

Functional connectivity for bivariate phase synchronization within the frequency bands showed significant differences between the CR stimulation and the control group. CR stimulation subjects showed a lower value than the control group corresponding to a greater decrease for all feature metrics derived for continuous signal phase synchronization in frequency bands. This denotes a greater decrease in functional segregation for the CR stimulation group. For the theta frequency band, two-sample tests showed increased modularity in the treatment group and a decrease in the control group. Transitivity was also significant between groups although it decreased in both groups, likely due to 4AP washout effects, which leads to more synchronized neuronal firing as concentration declines, similar to postictal seizure dynamics. Studies have shown that in some cases connectivity increases as seizures approach termination [87]. Binary connectivity matrices showed high significance for modularity, with an increase in the treatment group and a decrease in the control group, suggesting different brain-state responses between groups due to CR stimulation.

For the alpha frequency band weighted FC matrices did not return significant results in the permutation tests, but the binary connectivity matrices did show significant measures between CR stimulation and control groups, in particular, showing high significance for functional integration measures: shortest path and betweenness centrality. Other metrics such as degree, clustering coefficient and transitivity were also significant.

In the beta frequency band, significant results were observed for the weighted FC norm, maximum eigenvalue, and network measures of average degree, clustering coefficient, and transitivity. These metrics showed a general decrease in the treatment group compared to an increase in the control group, except for FC norm and transitivity, which also decreased in the control group. The significance of the norm and maximum eigenvalue confirms modulation in network phase coordination in the beta frequency band. The threshold sweep of binary FC matrices revealed that modularity and functional integration measures were also significant.

For the lower gamma frequency band, the average degree and clustering coefficient exhibited a significant relative percentage change, confirmed by TWFA. Lower gamma metrics showed a decrease in the treatment group and an increase in the control group, suggesting CR stimulation disrupts phase coordination in this frequency band. The reasons for increased connectivity in the control group remain unclear, but postictal activity may play a role in inducing clustered neuronal activity and increased connectivity as epileptiform activity terminates [87].

Threshold sweep analysis of binary FC matrices compensated for potential limitations in weighted FC tests, validating the results further. Notably, in the alpha band, functional integration metrics such as betweenness centrality and average degree showed high significance, with both groups exhibiting decreases. Similar findings in the beta band were confirmed for degree, shortest path length, and betweenness centrality.

Nodes with high betweenness centrality can act as epileptogenic hubs, promoting the spread of seizures. Targeting these hubs with interventions like electrical stimulation may help disrupt seizure propagation by modulating critical communication pathways [74].

In summary our results showed significant results for several phase synchronization bivariate features, including network measures of both functional segregation and integration for all frequency bands.

However, at a high level the most pronounced changes in functional segregation measures was shown by phase synchronization in the lower gamma, specifically the clustering coefficient. This was cross-validated with different statistical analyses (i.e. TWFA, relative percentage change, 2-sample permutation tests), consistently showing that for the lower gamma band the clustering coefficient in the CR stimulation group had a decrease in mean values compared to the control group which showed an increase (See Supplemental Tables and Figure 6), in alignment with previous research findings [87].

For functional integration network measures from FC matrices in the alpha band, shortest path and betweenness centrality were highly significant across all statistical analyses. As these metrics indicate a reduction in the efficiency of global information transfer, suggesting that CR stimulation disrupted long-range communication and enhanced the segregation of brain networks.

The beta, theta and gamma have a mix of significant metrics for functional integration and segregation, both share high significance for modularity across 2-sample tests and TWFA. The spike relative phase showed significant differences for all statistical tests for the maximum eigenvalue and also for modularity and the shortest path. This in general confirms that there was a larger induced desynchronization effect on the spikes between different channels in the CR stimulation group compared to the control group. As the CR stimulation group showed a larger change in overall relative spike phase. These frequency-specific effects show the potential of CR stimulation to disrupt epileptiform synchronization. The different behaviors motivate further research to analyze dynamic network measures to discover phases of ictogenesis that are particularly suitable to abort the seizure [88]. Based on our results obtained from the different frequency bands there might be different periods or phases of seizures and epileptiform activity that might increase treatment efficacy.

Other exploratory analyses that might provide relevant insights could be the study of cross-frequency and phase-amplitude interactions and their modulation with CR stimulation [89,90].

The technical limitations of this study consist of the set-up of stimulation contacts as it is possible that electrical stimulation from the different electrode contacts is still stimulating the same network (i.e. hippocampal formation). That is, there might be an overlap in network activation at each electrical stimulus despite its source being in different anatomical sites with a vertical separation of 1 mm. For long-lasting CR induced desynchronization to take place, stimulation contacts must not activate strongly overlapping subnetworks [55]. This could be the strongest limitation and the reason why not stronger results were observed (e.g. statistical significance in univariate power). The first step to further improve stimulation outcome will be an adequately adjusted placement of stimulation contacts according to theoretical and in vivo CR studies [20, 55, 91].

Another experimental limitation in this specific study is the use of 4-AP. We are attempting to measure modulation pre-post CR during the induction of sustained epileptiform activity through intracranial injection of 4-AP which blocks potassium channels [27]. So, as long as 4-AP is affecting the network, suppression of epileptiform activity will be partial. In this case we are comparing it against the control condition where no CR stimulation was delivered.

Regardless of the above limitations and of the experimental and technical setup being sub-optimal to test the full potential of CR stimulation effects, we nevertheless showed quantifiable changes between pre-CR and post-CR periods and between CR stimulation and control groups.

Additionally, we developed a comprehensive analytical framework with different statistical tools that evaluates the network properties within each period and their inter periods modulation through analysis using univariate and bivariate features extracted from spike-events and the LFP signal. The developed framework uses a comprehensive set of tests using different features and derived measurements including analysis in the spectral domain for power modulation and phase coordination. This framework is also useful for the analysis of modulation of EEG signals and will be used and applied to other datasets in the future.

Moreover, further clinical research would be instrumental to understanding the extent and the robustness in efficacy of CR stimulation for treatment of DRE. The methodology implemented in this study exemplifies the translational potential of CR stimulation, clinical applications in epilepsy management and a mathematical framework to analyze and evaluate acute changes caused by electrical stimulation. These results broaden our understanding of CR stimulation by implementing RVS-CR in an in vivo model of sustained epileptiform activity. Future research involving a comparative analysis against other electrical stimulation modalities like DBS, RNS, VNS, or transcranial Direct Current Stimulation (Fisher et al. 2023), might elucidate the relative advantages of CR stimulation.

In conclusion, the urgency for effective interventions to mitigate the debilitating impacts of epilepsy remains a challenge. This study highlights the potential of CR stimulation as a viable intervention to desynchronize epileptiform activity and improve seizure control in DRE.

## Acknowledgements

Daniel Ehrens received support for this research as a Howard Hughes Medical Institute Gilliam Fellow. Sridevi V. Sarma received support by NIH 1R21NS103113-01A1. Yitzhak Schiller received support from a Technion Israel Institute of Technology internal grant for collaboration with JHU faculty (R2026493). Peter A. Tass received support from the Vaughn Bryson fund. The first author wishes to recognize the exceptional diversity within this multidisciplinary team, which unifies a wide array of ethnic, cultural, spiritual, and ideological backgrounds. Diverse perspectives enrich the research process fostering innovative perspectives and strengthens our community and commitment to advancing epilepsy care and improving global health and quality of life for all individuals and their families.

