## Supplementary_Material for "Electrical Coordinated Reset Stimulation Induces Network Desynchronization in an in Vivo Model of Status Epilepticus"

### Supplemental Materials

### Figures

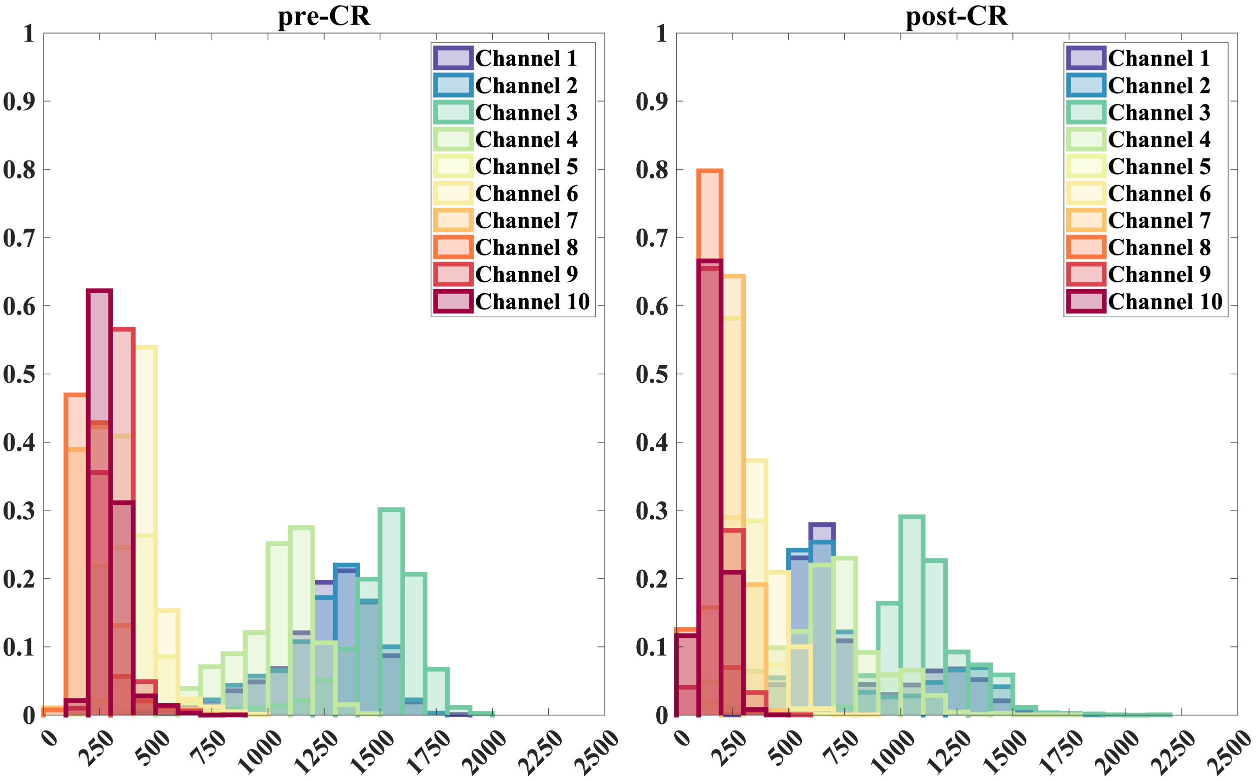


**Supplemental Figure 1.** Univariate spike amplitude analysis. Spike amplitude distribution for each channel is calculated for the different periods for subject CR-stim 1. Plot on the left shows the distribution of spike amplitude values for the pre-CR stimulation period. On the right plot, the distribution for spike amplitude during the post CR stimulation period is shown. Each color represents a different recording-site channel.

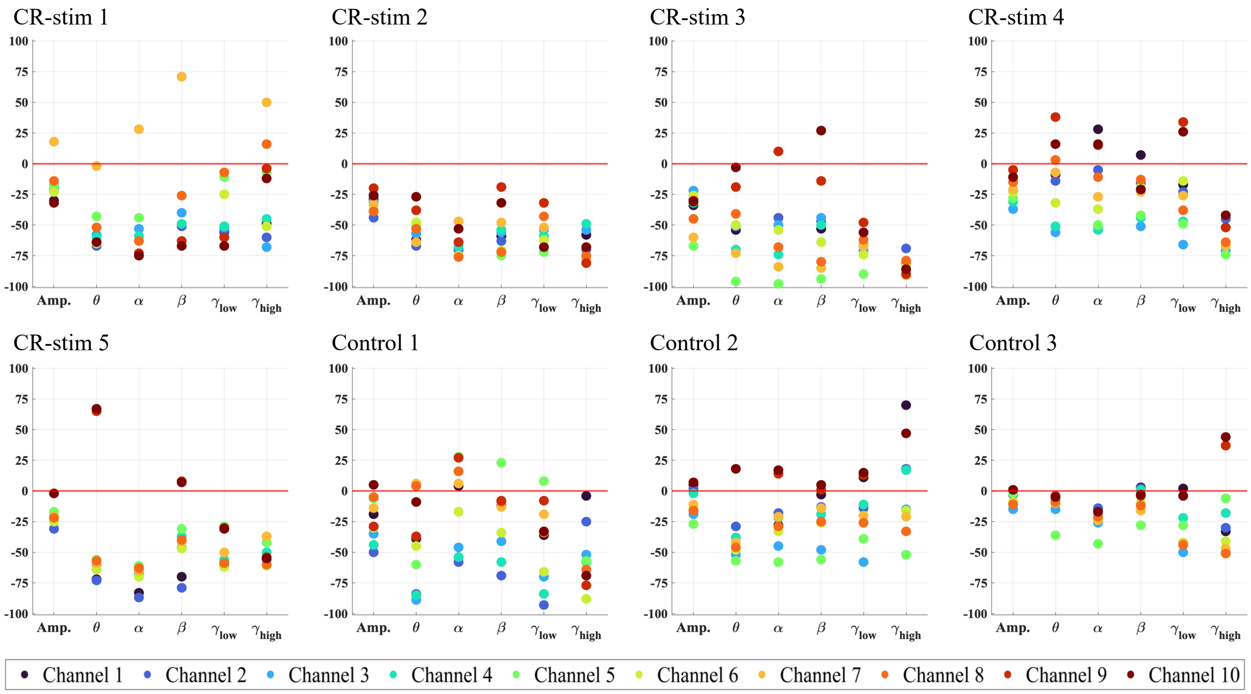


**Supplemental Figure 2.** Differential percentage per channel. CR stim group (N=5), and Control group (N=3) subjects feature-differential per channel. Each plot shows the mean differential percentage for each feature (amplitude and each frequency band) per subject. Each colored circle represents a different recording-site channel.

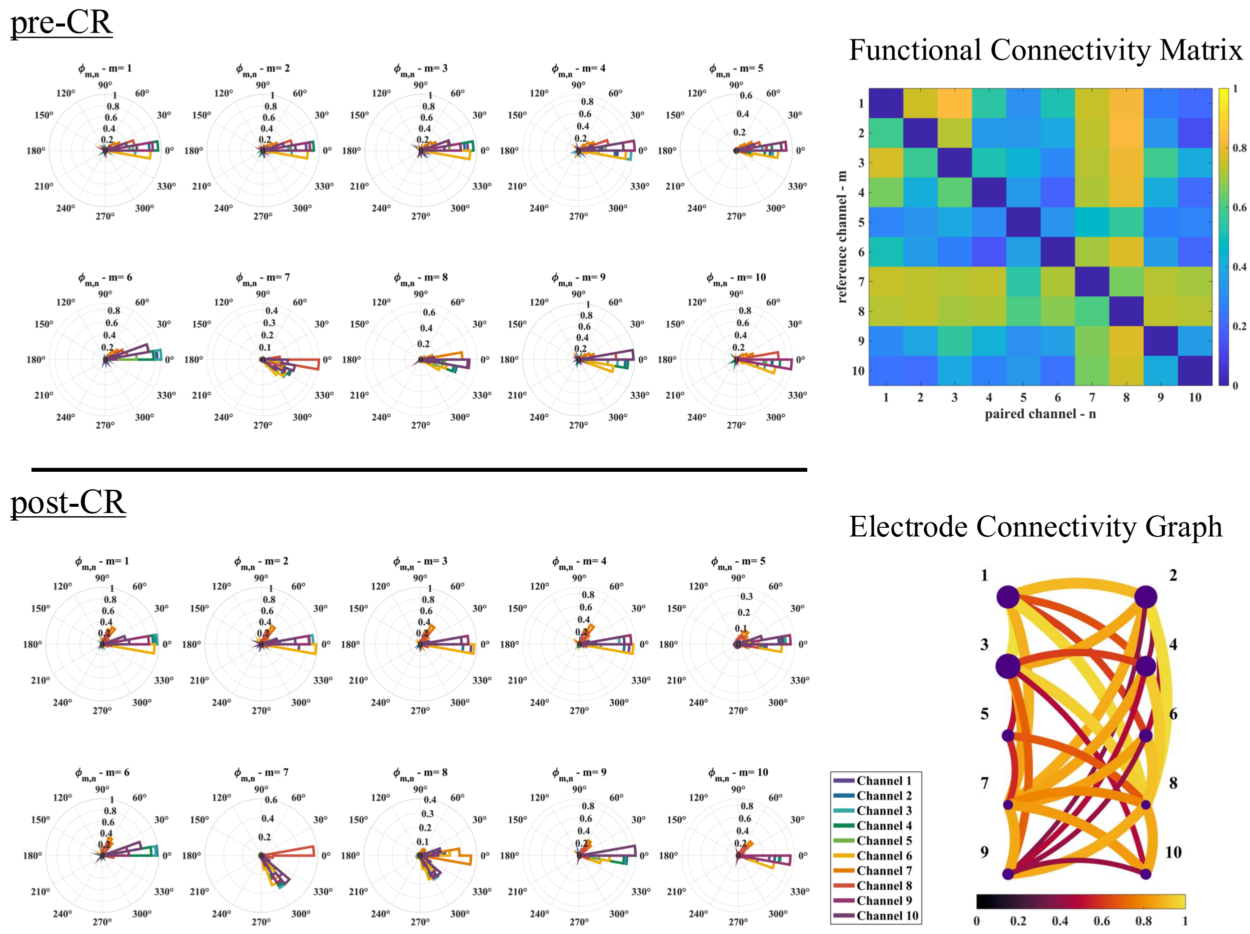


**Supplemental Figure 3.** Relative Spike Phase - bivariate analysis. Bivariate analysis for subject CR-stim 1. The top left panel shows the pre-CR bivariate relative spike phase polar histograms for all ten reference channels, *m,* in relation to the other channels. The bottom left panel shows the post-CR relative spike phase calculated for each channel. The panel on the right shows at the top a functional connectivity matrix constructed by performing the 2-sample Kuiper test on all channel pairs, displaying values between 0 and 1 describing the change between pre-CR and post-CR periods. In the bottom right plot, we show the corresponding electrode connectivity graph, where the diameter of the nodes corresponds to the spike amplitude and the edge width represents the strength in the connection given by the Kuiper statistic.

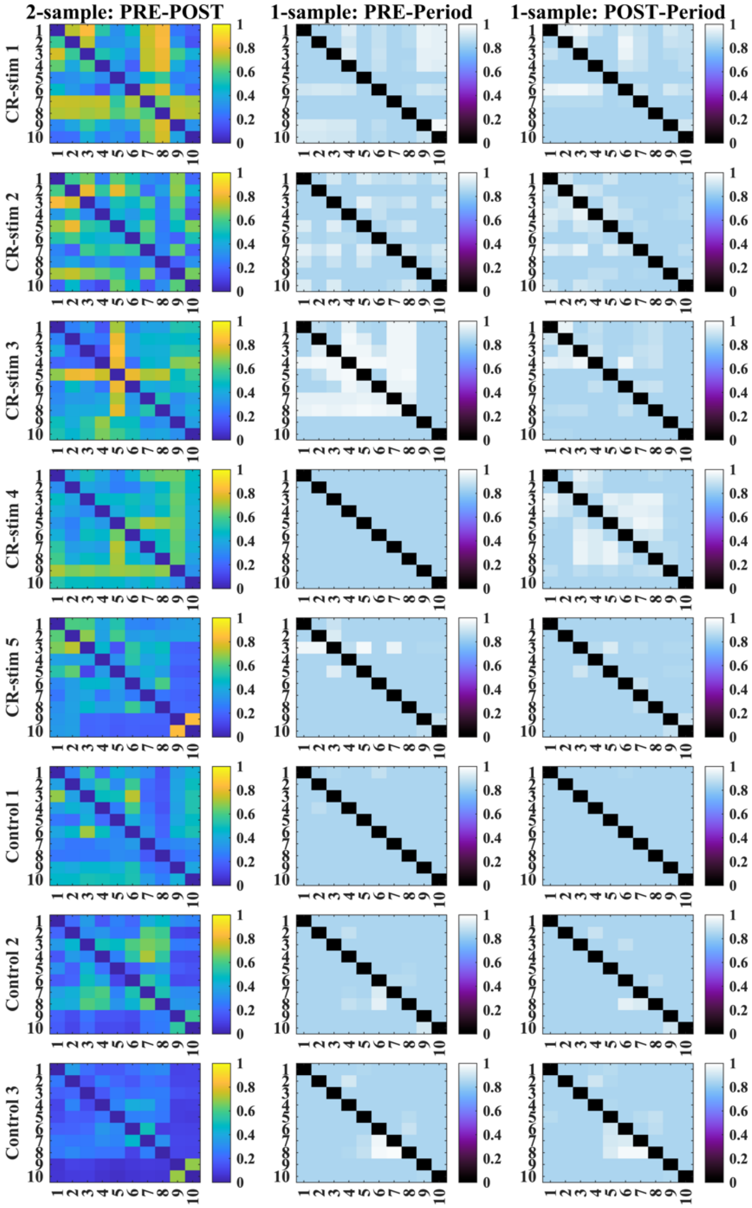


**Supplemental Figure 4.** Bivariate spike relative phase 2-sample and 1-sample Functional Connectivity (FC) matrices. FC matrices are calculated using the 2-sample Kuiper test for the first column (as seen in Figure 3), and the 1-sample Kuiper test for the pre-period in the center column and for the post-period in the third column. Kuiper tests were used to calculate phase coordination for each channel-pair's spike relative phase. The 2-sample Kuiper test was used to compare the pre-post-CR circular distributions and the 1-sample Kuiper test evaluated synchronization as it compared the observed distribution to a uniform circular distribution. The FC matrices are shown for all subjects (rows).

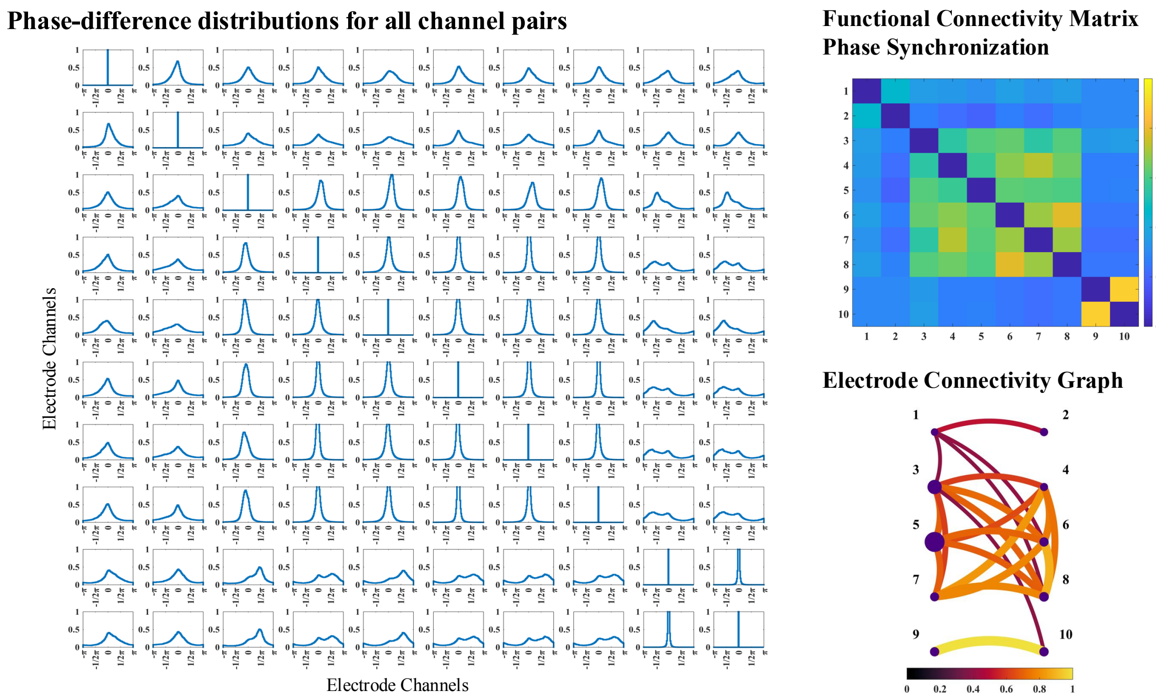


**Supplemental Figure 5.** Phase difference distribution and FC matrix construction. Bivariate analysis for subject CR-stim 5, beta band. On the left a grid of ten by ten contains the phase-difference distributions for each channel pair. On the right the functional connectivity matrix is created using the 1-sample Kuiper test from each phase-difference distribution to calculate phase synchronization for each channel pair. In the bottom right plot, we show the corresponding electrode connectivity graph, where the diameter of the nodes corresponds to the univariate power in the beta band for each corresponding channel, and the edge width represents the strength in phase synchronization between nodes given by the Kuiper statistic.

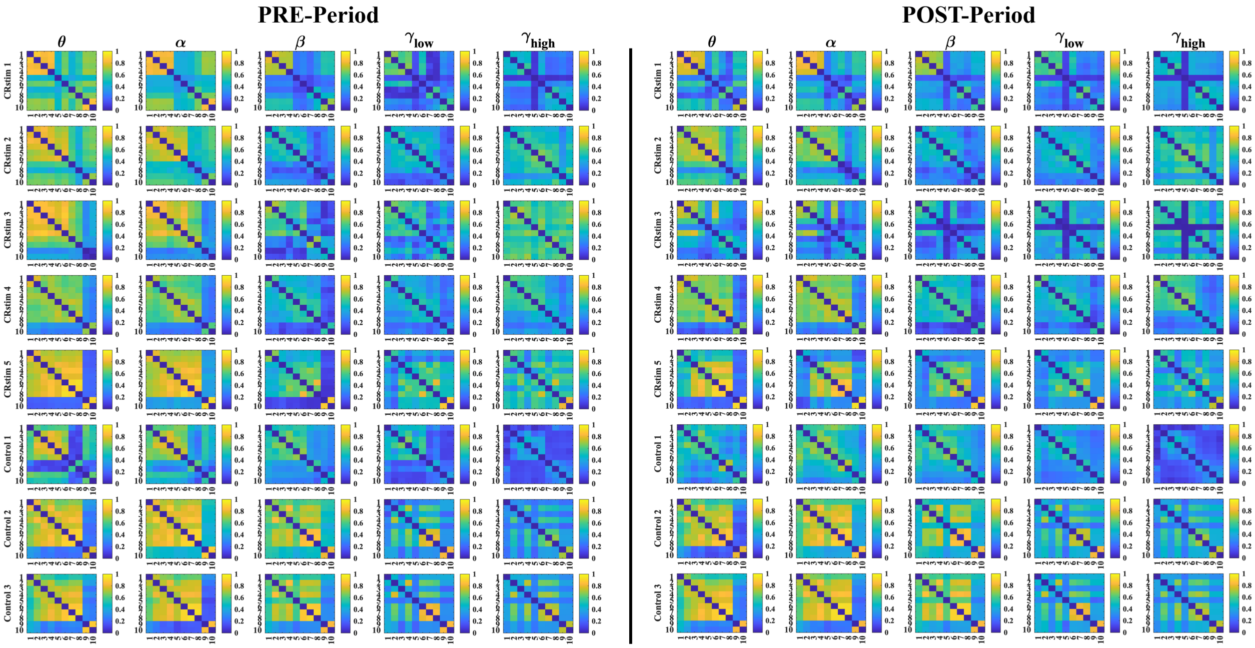


**Supplemental Figure 6.** Bivariate phase coordination 1-sample Functional Connectivity (FC) matrices. FC matrices are calculated using the 1-sample Kuiper test to calculate phase coordination for each channel-pair's phase-difference distribution.

On the left the pre-CR or pre-no-stim period FC matrices are shown for all subjects (rows) and all frequency bands (columns). Similarly on the right we show the post-CR or post-no-stim period FC matrices for all subjects and all frequencies.

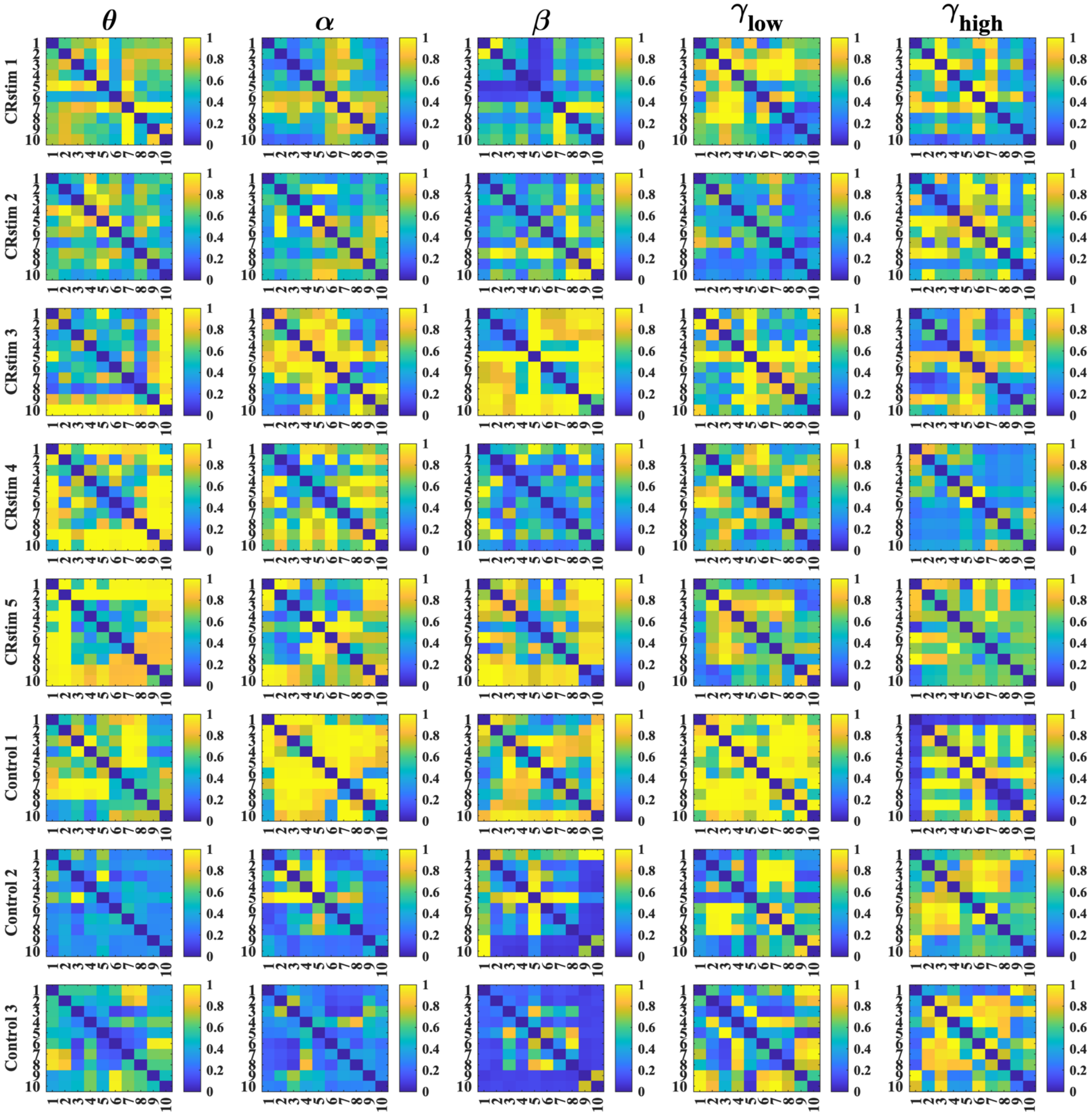


**Supplemental Figure 7.** Bivariate phase coordination 2-sample Functional Connectivity (FC) matrices.
FC matrices are calculated using the 2-sample Kuiper test to calculate phase coordination for each channel-pair's phase-difference distribution.
2-sample FC matrices are calculated using both the pre-CR or pre-no-stim period and the post-CR or post-no-stim period .FC matrices are shown for all subjects (rows) and all frequency bands (columns).

### Statistical Results Tables

**Supplemental Table 1**

| Statistical test analysis and network feature | *P* | Cohen's D |
|---|---|---|
| Spk Rel Phi 2-sample Norm | 0.0194 | 2.31 |
| Spk Rel Phi 2-sample Max Eig | 0.0271 | 2.14 |
| Spk Rel Phi 2-sample Avg Degree | 0.0368 | 2.26 |
| Spk Rel Phi 2-sample Clust Coeff | 0.0356 | 2.26 |
| Spk Rel Phi 2-sample Transitivity | 0.0362 | 1.93 |
| Spk Rel Phi Two-way Factorial Max Eig (Treatment) | 0.0376 | 1.41 |

**Supplemental Table 1.** Statistically significant results, on spike relative phase for all network features. Different statistical test analyses and network features are shown for the weighted Functional Connectivity matrix from which network graph measures are derived and tested for significance using different permutation tests analyses.

**Supplemental Table 2**

| Statistical test analysis and network feature | *P* | Cohen's D |
|---|---|---|
| 2-sample Modularity - Theta | 0.03 | -2.28 |
| Rel %Delta Avg Degree - Gamma-low | 0.0428 | -1.84 |
| Two-way Factorial Norm (Treatment) - Beta | 0.0368 | -1.99 |
| Two-way Factorial Max Eig (Treatment) - Beta | 0.0194 | -2.15 |
| Two-way Factorial Max Eig (Interaction) - Gamma-low | 0.0325 | -1.91 |
| Two-way Factorial Avg Degree (Treatment) - Beta | 0.0103 | -2.35 |
| Two-way Factorial Avg Degree (Interaction) - Gamma-low | 0.037 | -2.12 |
| Two-way Factorial Clust Coeff (Treatment) - Beta | 0.0134 | -2.35 |
| Two-way Factorial Clust Coeff (Interaction) - Gamma-low | 0.0381 | -2.12 |
| Two-way Factorial Transitivity (Treatment) - Beta | 0.0368 | -1.80 |
| Two-way Factorial Transitivity (Time) - Theta | 0.0235 | -1.31 |

**Supplemental Table 2.** Statistically significant results, on phase synchronization features for all frequencies. Different statistical test analyses and network features are shown for the weighted Functional Connectivity matrix from which network graph measures are derived and tested for significance using multiple permutation tests.

**Supplemental Table 3**

| Statistical test analysis and network feature | *P* | Cohen's D |
|---|---|---|
| Spk Rel Phi 2-sample Avg Degree - FC Thresh 0.10 | 0.037 | 1.07 |
| Spk Rel Phi 2-sample Avg Degree - FC Thresh 0.20 | 0.0411 | 1.52 |
| Spk Rel Phi 2-sample Avg Degree - FC Thresh 0.25 | 0.0207 | 1.74 |
| Spk Rel Phi 2-sample Avg Degree - FC Thresh 0.30 | 0.0117 | 2.09 |

| | | |
|---|---|---|
| Spk Rel Phi 2-sample Avg Degree - FC Thresh 0.35 | 0.0194 | 2.17 |
| Spk Rel Phi 2-sample Clust Coeff - FC Thresh 0.30 | 0.0365 | 1.73 |
| Spk Rel Phi 2-sample Clust Coeff - FC Thresh 0.35 | 0.0365 | 1.64 |
| Spk Rel Phi 2-sample Clust Coeff - FC Thresh 0.45 | 0.0382 | 1.87 |
| Spk Rel Phi 2-sample Modularity - FC Thresh 0.15 | 0.042 | -1.30 |
| Spk Rel Phi 2-sample Shortest Path - FC Thresh 0.15 | 0.0407 | -1.30 |
| Spk Rel Phi 2-sample Shortest Path - FC Thresh 0.60 | 0.0272 | 2.41 |
| Spk Rel Phi 2-sample Shortest Path - FC Thresh 0.65 | 0.0362 | 2.41 |
| Spk Rel Phi 2-sample Transitivity - FC Thresh 0.25 | 0.0368 | 1.99 |
| Spk Rel Phi Two-way Factorial Betweenness (Treatment) - FC Thresh 0.90 | 0.0212 | 1.53 |
| Spk Rel Phi Two-way Factorial Modularity (Treatment) - FC Thresh 0.90 | 0.0212 | 0.00 |
| Spk Rel Phi Two-way Factorial Shortest Path (Treatment) - FC Thresh 0.90 | 0.0212 | 1.77 |

**Supplemental Table 3.** Statistically significant results, on bivariate spike relative phase. Different statistical test analyses and network features are shown for the threshold sweep, in order to calculate an adjacency matrix, and compute network graph measures.

**Supplemental Table 4**

| Statistical test analysis and network feature | *P* | Effect size |
|---|---|---|
| 2-sample Betweenness - FC Thresh 0.55-Gamma-low | 0.0364 | 3.03 |
| 2-sample Betweenness - FC Thresh 0.60-Gamma-low | 0.0365 | 2.37 |
| 2-sample Betweenness - FC Thresh 0.65-Alpha | 0.0194 | 2.80 |
| 2-sample Betweenness - FC Thresh 0.70-Alpha | 0.0097 | 2.31 |
| 2-sample Betweenness - FC Thresh 0.75-Alpha | 0.0194 | 2.01 |
| 2-sample Clust Coeff - FC Thresh 0.35-Gamma-low | 0.049 | -2.13 |
| 2-sample Modularity - FC Thresh 0.20-Beta | 0.0448 | -1.42 |
| 2-sample Modularity - FC Thresh 0.75-Alpha | 0.0194 | 2.09 |
| 2-sample Shortest Path - FC Thresh 0.55-Gamma-low | 0.0358 | 1.95 |
| 2-sample Shortest Path - FC Thresh 0.60-Gamma-low | 0.0493 | 2.14 |
| 2-sample Shortest Path - FC Thresh 0.65-Alpha | 0.0391 | 1.68 |
| 2-sample Shortest Path - FC Thresh 0.75-Alpha | 0.0194 | 1.68 |
| 2-sample Transitivity - FC Thresh 0.60-Gamma-low | 0.0369 | -1.68 |
| Rel %Delta Avg Degree - FC Thresh 0.20-Alpha | 0.0046 | -1.97 |
| Rel %Delta Betweenness - FC Thresh 0.15-Alpha | 0.0368 | 2.97 |
| Rel %Delta Betweenness - FC Thresh 0.20-Alpha | 0.0066 | 5.29 |
| Rel %Delta Betweenness - FC Thresh 0.40-Gamma-low | 0.0135 | -1.73 |
| Rel %Delta Betweenness - FC Thresh 0.60-Alpha | 0.0358 | -2.97 |
| Rel %Delta Clust Coeff - FC Thresh 0.15-Alpha | 0.0493 | -2.59 |

| | | |
|---|---|---|
| Rel %Delta Clust Coeff - FC Thresh 0.20-Alpha | 0.039 | -2.72 |
| Rel %Delta Clust Coeff - FC Thresh 0.20-Gamma-low | 0.0194 | -2.95 |
| Rel %Delta Modularity - FC Thresh 0.10-Gamma-low | 0.0376 | 2.05 |
| Rel %Delta Modularity - FC Thresh 0.20-Alpha | 0.0194 | 1.77 |
| Rel %Delta Modularity - FC Thresh 0.20-Gamma-high | 0.0377 | 1.86 |
| Rel %Delta Shortest Path - FC Thresh 0.15-Alpha | 0.0356 | 2.51 |
| Rel %Delta Shortest Path - FC Thresh 0.20-Alpha | 0.0194 | 2.12 |
| Rel %Delta Shortest Path - FC Thresh 0.55-Alpha | 0.0354 | -1.89 |
| Rel %Delta Shortest Path - FC Thresh 0.60-Alpha | 0.0194 | -2.23 |
| Rel %Delta Transitivity - FC Thresh 0.15-Alpha | 0.0368 | -2.54 |
| Rel %Delta Transitivity - FC Thresh 0.20-Alpha | 0.039 | -2.45 |
| Rel %Delta Transitivity - FC Thresh 0.60-Alpha | 0.0101 | 1.63 |
| Two-way %Delta Avg Degree (Interaction) - FC Thresh 0.15-Alpha | 0.0363 | -2.65 |
| Two-way %Delta Avg Degree (Interaction) - FC Thresh 0.20-Alpha | 0.039 | -1.87 |
| Two-way %Delta Avg Degree (Treatment) - FC Thresh 0.10-Beta | 0.0477 | -1.22 |
| Two-way %Delta Avg Degree (Treatment) - FC Thresh 0.15-Beta | 0.0346 | -1.79 |
| Two-way %Delta Avg Degree (Treatment) - FC Thresh 0.20-Beta | 0.0194 | -2.51 |
| Two-way %Delta Avg Degree (Treatment) - FC Thresh 0.25-Beta | 0.0051 | -5.27 |
| Two-way %Delta Avg Degree (Treatment) - FC Thresh 0.30-Beta | 0.0194 | -4.00 |
| Two-way %Delta Avg Degree (Treatment) - FC Thresh 0.35-Beta | 0.0194 | -2.19 |
| Two-way %Delta Avg Degree (Treatment) - FC Thresh 0.40-Beta | 0.0498 | -1.56 |
| Two-way %Delta Avg Degree (Treatment) - FC Thresh 0.50-Beta | 0.0368 | -1.74 |
| Two-way %Delta Avg Degree (Treatment) - FC Thresh 0.55-Beta | 0.0363 | -1.63 |
| Two-way %Delta Betweenness (Interaction) - FC Thresh 0.10-Alpha | 0.0391 | 1.15 |
| Two-way %Delta Betweenness (Interaction) - FC Thresh 0.15-Alpha | 0.0391 | 2.65 |
| Two-way %Delta Betweenness (Interaction) - FC Thresh 0.20-Alpha | 0.039 | 1.87 |
| Two-way %Delta Betweenness (Treatment) - FC Thresh 0.15-Beta | 0.0049 | 1.88 |
| Two-way %Delta Betweenness (Treatment) - FC Thresh 0.20-Beta | 0.0391 | 1.50 |
| Two-way %Delta Betweenness (Treatment) - FC Thresh 0.50-Gamma-high | 0.036 | 1.63 |
| Two-way %Delta Betweenness (Treatment) - FC Thresh 0.60-Beta | 0.0194 | -1.52 |
| Two-way %Delta Clust Coeff (Interaction) - FC Thresh 0.20-Alpha | 0.039 | -2.87 |
| Two-way %Delta Clust Coeff (Interaction) - FC Thresh 0.20-Gamma-low | 0.0182 | -3.02 |
| Two-way %Delta Clust Coeff (Interaction) - FC Thresh 0.45-Alpha | 0.0445 | -2.45 |
| Two-way %Delta Clust Coeff (Time) - FC Thresh 0.30-Alpha | 0.0356 | -0.89 |
| Two-way %Delta Clust Coeff (Treatment) - FC Thresh 0.20-Beta | 0.0194 | -2.24 |

| | | |
|---|---|---|
| Two-way %Delta Clust Coeff (Treatment) - FC Thresh 0.25-Beta | 0.0194 | -5.68 |
| Two-way %Delta Clust Coeff (Treatment) - FC Thresh 0.30-Beta | 0.0194 | -3.05 |
| Two-way %Delta Clust Coeff (Treatment) - FC Thresh 0.45-Gamma-high | 0.0383 | 1.41 |
| Two-way %Delta Modularity (Time) - FC Thresh 0.45-Beta | 0.0295 | 0.54 |
| Two-way %Delta Modularity (Treatment) - FC Thresh 0.15-Beta | 0.045 | 0.94 |
| Two-way %Delta Modularity (Treatment) - FC Thresh 0.20-Beta | 0.0409 | 1.60 |
| Two-way %Delta Modularity (Treatment) - FC Thresh 0.25-Beta | 0.0194 | 2.37 |
| Two-way %Delta Modularity (Treatment) - FC Thresh 0.35-Beta | 0.024 | 1.15 |
| Two-way %Delta Norm (Treatment) - FC Thresh 0.10-Beta | 0.0368 | -1.99 |
| Two-way %Delta Shortest Path (Interaction) - FC Thresh 0.20-Alpha | 0.039 | 1.87 |
| Two-way %Delta Shortest Path (Interaction) - FC Thresh 0.50-Gamma-low | 0.0481 | -2.43 |
| Two-way %Delta Shortest Path (Interaction) - FC Thresh 0.55-Alpha | 0.0252 | -1.71 |
| Two-way %Delta Shortest Path (Interaction) - FC Thresh 0.60-Alpha | 0.0313 | -1.70 |
| Two-way %Delta Shortest Path (Time) - FC Thresh 0.25-Gamma-high | 0.0121 | -0.47 |
| Two-way %Delta Shortest Path (Treatment) - FC Thresh 0.15-Beta | 0.007 | 1.88 |
| Two-way %Delta Shortest Path (Treatment) - FC Thresh 0.50-Gamma-high | 0.0383 | 1.85 |
| Two-way %Delta Shortest Path (Treatment) - FC Thresh 0.60-Beta | 0.0194 | -1.88 |
| Two-way %Delta Transitivity (Interaction) - FC Thresh 0.15-Alpha | 0.0485 | -2.71 |
| Two-way %Delta Transitivity (Interaction) - FC Thresh 0.20-Alpha | 0.039 | -2.52 |
| Two-way %Delta Transitivity (Interaction) - FC Thresh 0.30-Alpha | 0.0475 | -1.72 |
| Two-way %Delta Transitivity (Interaction) - FC Thresh 0.30-Beta | 0.0452 | -2.45 |
| Two-way %Delta Transitivity (Treatment) - FC Thresh 0.15-Beta | 0.035 | -1.98 |

**Supplemental Table 4.** Statistically significant results, on phase synchronization features for all frequencies. Different statistical test analyses and network features are shown for the threshold sweep, in order to calculate an adjacency matrix and compute network graph measures.
